# Comparison of tick surveillance approaches: Utilizing trailhead tick-check stations to support tick surveillance and community education

**DOI:** 10.64898/2026.06.01.728881

**Authors:** Sabrina Gobran, Caroline Fagan, Lawson Dawe, Emma K. Harris, Christopher M Roundy, Shelby Cagle, Nicole Kelp, Karla-Saavedra Rodriguez, Elizabeth Hemming-Schroeder

## Abstract

**Introduction:** With rising tick-borne disease (TBD) cases and the geographical expansion of tick populations, the need for effective surveillance and public education regarding local risk is crucial. This study assessed the effectiveness of tick-check stations as a tool for tick surveillance, and their impact on community knowledge, attitude, and practices (KAP) related to ticks and TBD.

**Methods:** To assess the effectiveness of tick-check stations for surveillance, we evaluated station engagement and compared tick density estimates, species composition, and life-stage distributions with those obtained through concurrent active surveillance. In addition, we compared submission numbers and tick species and life stages to those collected through a mail-in submission system conducted by the Colorado Department of Public Health and Environment (CDPHE). To quantify feasibility, we estimated effort per tick and compared effort across simulated sampling scenarios. Finally, in-person surveys were conducted at trailheads to assess baseline tick KAP and to evaluate differences between sites with and without tick-check stations

**Results:** Engagement with tick-check stations was sustained throughout the study. Temporal tick densities estimated from tick-check station submissions were correlated with density estimates from active surveillance (R = 0.534), and species composition and life-stage distributions did not significantly differ between methods. Tick-check stations required less effort per tick than active surveillance when sampling sites were nearby or tick densities were low, whereas sites that were farther away or had higher tick densities required less effort per tick under a hybrid surveillance approach. When asked to list tick-borne pathogens in Colorado, 47% of survey participants who had read tick-check station signage identified Rocky Mountain spotted fever compared with 20% of participants in the control group (p = 0.007; odds ratio). Notably, a low proportion of survey participants (24%) reported performing tick-checks to prevent tick bites.

**Conclusion:** Tick-check stations can provide tick density estimates comparable to active surveillance while requiring less effort in many scenarios, particularly in low-density settings. Our findings also highlight opportunities for targeted outreach to address gaps in TBD knowledge. As both a surveillance and educational tool, tick-check stations offer a sustainable approach for expanding tick monitoring in resource-limited settings.

## 1 Introduction

Since 2004, the number of reported tick-borne disease (TBD) cases in the United States has been steadily rising (1). Concurrently, the geographical distribution of TBD incidence has expanded, mirroring the expansion of tick vector populations (2,3). Multiple factors contribute to these trends, including shifts in habitat suitability induced by climate change, changes in land-use affecting host availability, increased awareness and reporting of TBDs, and the implementation of more tick surveillance systems and methods (4,5). Given the persistence of climate change and environmental disturbances, ticks and their associated pathogens are likely to undergo continued shifts in distribution (6–8). Therefore, effective surveillance systems will be critical for the early detection of invasive vectors and emerging pathogens, ultimately supporting effective control and prevention of TBDs (4,9–11).

While tick surveillance is essential for understanding and mitigating disease risk, its incorporation into routine public health programming is often constrained by cost, especially in areas with low TBD incidence (5). Active surveillance, typically involving systematic drag sampling over vegetation to collect questing ticks, provides geographically accurate, standardized estimates of tick presence and density (10,12). However, this approach is resource-intensive, requiring significant time, labor, and logistical investment (13,14). Alternatively, passive surveillance programs—typically relying on tick submissions from community members—are less cost-prohibitive, but yield less standardized and spatially accurate data, and are often biased by level of community awareness and engagement (9,14,15).

Current tick surveillance objectives, at both state and local agencies, function to detect the presence and abundance of ticks and evaluate tick-borne pathogens to understand public health risks (5). Most surveyed agencies conducted tick surveillance through passive programs (typically via a mail-in system), or by irregular active surveillance activities. However, up to 35% of these agencies reported being unable to carry out the main objectives of their tick surveillance programs, primarily due to lack of funding and resources. Although passive surveillance programs may be limited in standardization and spatial resolution, this approach can still help address certain public health objectives when systematic active surveillance is not feasible. Passive surveillance may even prove superior to active sampling in measuring realized tick-human encounters, a dimension not typically captured by most active surveillance methodologies (16).

Given the limitations of traditional passive surveillance approaches, namely, the underutilization of these programs by the public, that data may be biased and lack standardization, and the high costs and labor demands of active surveillance, we developed an alternative method based on implementation of trailhead tick-check stations. These stations provide a low-effort, accessible platform for academic researchers and public health agencies to collect ticks by engaging community members to directly submit ticks encountered during outdoor activities at conveniently located submission stations. This approach leverages human-tick interactions while minimizing logistical burden. We hypothesized that trailhead tick-check stations could successfully capture geographically relevant data and provide estimates of tick density comparable to active surveillance while requiring fewer resources compared to traditional active surveillance. To evaluate this approach, we installed tick-check stations at four locations in Northern Colorado during spring-summer 2024 to coincide with the peak host-seeking activity season of *Dermacentor andersoni* and *Dermacentor variabilis* ticks. Trailheads were targeted for placement of tick-check stations, as tick encounters in Colorado are most likely to occur while recreating in natural areas (17).

Given that Colorado hikers and outdoor recreationalists are at a higher risk of encountering ticks and tick-borne pathogens (18,19), tick-check stations present a unique opportunity not only for surveillance but also for targeted public health outreach. The knowledge, attitudes, and practices (KAP) framework provides a structured approach to examine whether changes in knowledge and risk perception translate into sustained protective practices (20,21). Within this framework, knowledge refers to factual awareness of risks; attitudes encompass perceived risks, benefits, and barriers; and practices represent the behaviors that ultimately determine exposure and are shaped by both knowledge and attitudes. We hypothesized that informational signage provided at trailhead tick-check stations could enhance community KAP regarding ticks and TBD.

Previous studies into tick KAP have primarily focused on individuals with high occupational exposure to ticks (e.g., Forest Service employees, farmers, veterinarians) (22–24) or individuals with recreational exposure (25,26). Although outdoor workers often reported receiving inadequate information on tick bite prevention (23,27,28), these groups may be more accessible for intervention through structured training and resource provision. In contrast, individuals engaging in outdoor activities for recreation may be more difficult to target, as their risk varies widely with the frequency, duration, and type of activity (25). We conducted in-person surveys at two trailheads equipped with tick-check stations and at two control trailheads in Larimer County, Colorado to evaluate whether tick-check stations could serve as an effective education tool to target individuals with recreational exposure. Tick-check stations could be placed at trails with high tick exposure risk to encourage community implementation of tick-bite prevention strategies, ultimately promoting public health in natural areas.

## 2 Materials and methods

### 2.1 Study Design

The purpose of this study was to test the effectiveness of tick-check stations for passive tick surveillance and community education. To evaluate the effectiveness of tick-check stations against other surveillance methods, we compared our approach to traditional active sampling and a mail-in based tick submission program led by the Colorado Department of Public Health and Environment (CDPHE). To evaluate whether educational signage was effective at improving hiker KAP towards ticks and TBDs, we conducted in-person surveys at 2 trailheads with tick-check stations and 2 trailheads without (Figure 1).

**Figure 1.**
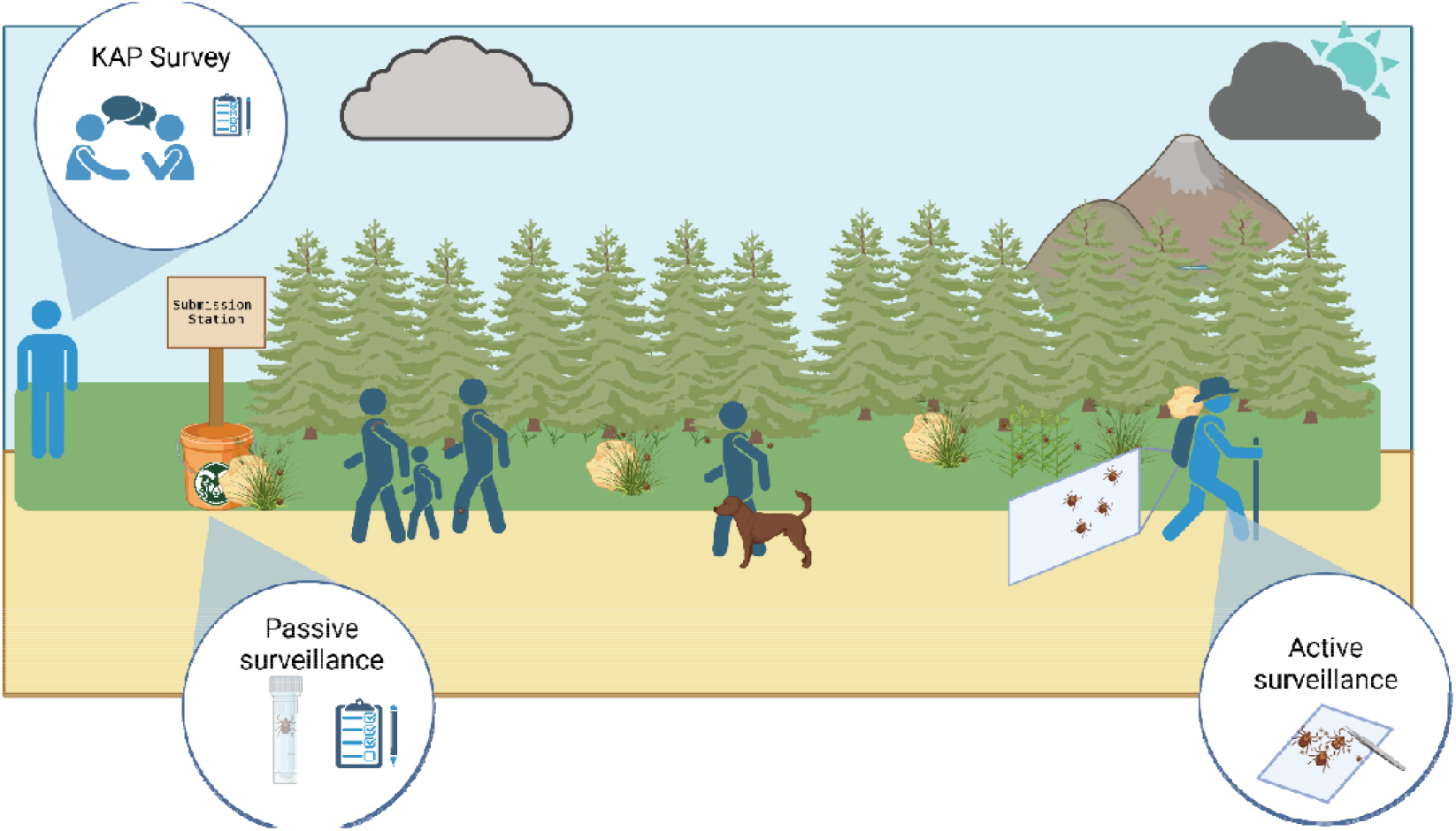
Study design.

### 2.2 Study Location

Study activities took place at six locations across Boulder and Larimer counties, Colorado (Table 1). Sites were located at both lower elevations (< 2000 m) in the foothills and higher elevations (> 2000 m) in the mountains to target *D. variabilis* and *D. andersoni*, respectively (17,29–31). All study activities (i.e., active sampling, tick-check station, KAP surveys) were conducted at two of our study sites (ELC, Soderberg). Active sampling and tick-check stations were conducted at two additional sites (Mud Lake, CSU Mountain Campus). To assess community KAP without the trailhead intervention, KAP surveys alone were delivered at Horsetooth Mountain and Spring Canyon trail. All sites were located on publicly accessible trails, except for the CSU Mountain Campus. While the CSU Mountain Campus is not a public natural area or trail system, this campus is populated with students in the summer and has a high tick encounter rate (32). The tick-check station was placed in a central location where it was accessible to the students.

**Table 1.** Site information.

| Site name | Location (lat, long) | Study activities | Land manager | County | Elevation (m) | Predicted tick species |
| --- | --- | --- | --- | --- | --- | --- |
| CSU Environmental Learning Center (ELC) | 40.55693, -105.01921 | Active sampling, tick-check station, KAP surveys | Colorado State University | Larimer | 1489 | *D. variabilis* |
| Soderberg | 40.53203,<br>-105.163 | Active<br>sampling,<br>tick-check<br>station,<br>KAP<br>surveys | Larimer<br>County | Larimer | 1664 | *D. variabilis* |
| CSU<br>Mountain<br>Campus | 40.56725,<br>-105.594 | Active<br>sampling,<br>tick-check<br>station | Colorado<br>State<br>University | Larimer | 2747 | *D.*<br>*andersoni* |
| Mud Lake | 39.97921,<br>-105.50823 | Active<br>sampling,<br>tick check<br>station | Boulder<br>County | Boulder | 2546 | *D.*<br>*andersoni* |
| Horsetooth<br>Mountain | 40.52353, -<br>105.18083 | KAP<br>surveys | Larimer<br>County | Larimer | 1764 | *D. variabilis* |
| Spring<br>Canyon trail | 40.52387, -<br>105.16530 | KAP<br>surveys | City of<br>Fort<br>Collins | Larimer | 1659 | *D. variabilis* |

### 2.3 Tick-Check Stations

Tick-check stations were designed to be placed at trailheads, allowing hikers to view information when entering and exiting the trail (Figure 2). At the CSU Mountain Campus, the tick-check station was centrally located to maximize participation. The front of the sign, which was viewed by hikers when approaching the trail, included information about ticks and TBDs in Colorado. The rear of the sign, which was viewed by hikers when exiting the trail, included information about how to conduct a tick check and participate in the tick submission program. Surveys and tick submission kits (including a survey, submission tube, disposable forceps, and disinfecting wipe) were provided in dispensers at the station. Hikers were directed to place completed submissions in a locked drop box attached to the station. Hikers were encouraged to submit a survey whether or not they found a tick during their tick check to quantify overall trail visitation. Stations were visited every 1-2 weeks to replenish supplies and collect submissions. Submitted ticks were transported to the lab alive for microscopic identification and then preserved at – 80°C. Materials and instructions for building the tick-check stations and survey questions are provided in the supplement (Supplementary Materials 1).

**Figure 2.**
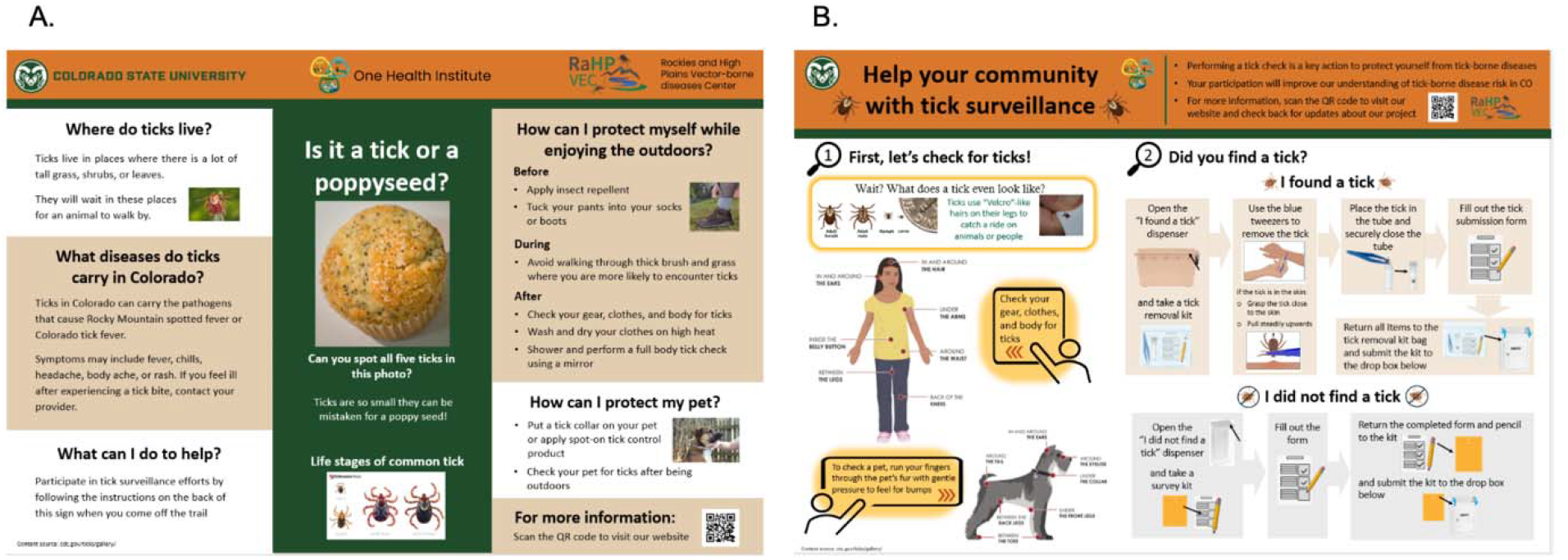
Trailhead sign front (A) and back (B).

### 2.4 Passive Surveillance: Mail-In System

Ticks were submitted to CDPHE by members of the public and veterinarians. Prior to submission, contributors completed a standardized form capturing key metadata, including the county of collection, specific geographic location (e.g., trail, park, or address), host associated (if applicable), and host type. Submitted ticks were double bagged in zip top bags and mailed to CDPHE for processing. Upon receipt, ticks were identified by state entomologists to species, life stage, sex, and engorgement status using microscopy and standard dichotomous keys. Results were reported back to the submitter.

### 2.5 Active Surveillance

We used a standard dragging approach for active surveillance, which involves dragging a 1-m^2^ white cloth along the forest floor and removing any attached ticks (33). Sampling was conducted along 20-meter transects parallel to the trail edge every 1-2 weeks. Drags were inspected every 5 m, and any attached ticks were removed with forceps and placed into uniquely identified vials. Transects were sampled on either side of the trail at 0 m, 5 m, and 10 m from the trail edge, where accessible. Starting points for each transect were randomly selected along the trail during each visit and data was entered electronically using the ODK Collect mobile data collection platform (https://getodk.org/). Ticks were transported back to the lab alive for microscopic identification, then stored at –80°C for long term storage.

### 2.6 Tick Identification

Ticks were kept alive at 4°C until they could be identified. Tick species and life stage were identified by microscopy and standard dichotomous keys (34). Briefly, life stage was established by determining the presence/absence of genital aperture (present in adults, absent in immatures) and number of legs. Tick species was identified by the size and number of goblet cells on the spiracular plates. Tick morphology was compared with the newly described *Dermacentor similis*, and we did not observe them in our area.

### 2.7 KAP Surveys

We designed a nine question, in-person, verbal survey consisting of Likert scale, Y/N, multiple choice, and open response questions to evaluate hiker KAP (Supplementary Materials 2) based on the design of Cuadera, *et al*. 2023 (35). Surveys were conducted in-person from late June to early September 2024. Survey respondents were approached and gave verbal consent to participate in the study as they were leaving the trail. Researchers read survey questions to the participant(s), and input participant verbal responses into the ODK platform. Responses were recorded without identifying information, although demographic information was collected (Supplementary Materials 3).

To assess knowledge, survey participants were asked to correctly identify 1) proper tick removal technique, 2) ticks from a series of images, and 3) TBDs endemic to Colorado. To assess attitudes, survey participants were asked to rate their concern for tick bites and TBDs on a scale of 1-5. To assess practices, survey participants were asked whether they take any precautions to prevent tick bites. Survey responses in the knowledge category were classified as incorrect or correct. Following guidelines published by the CDC, correct tick removal was defined as using tweezers and/or removing the tick by the head (36); any other response was considered incorrect for analysis purposes. Correct TBD identification was defined as Rocky Mountain spotted fever (RMSF) or Colorado tick fever (CTF) (37–40); any other responses, such as identifying Lyme disease, were classified as incorrect for analysis purposes. Open ended prevention practices were broadly classified as preventative or post-exposure behaviors. Preventative behaviors were subclassified as related to clothing (e.g. tucking pants into socks), repellent (e.g. applying DEET), and trail behavior (e.g. staying on designated trails). Post-exposure behaviors were subclassified as performing tick checks, taking a shower, or washing clothes after hiking.

### 2.8 Statistical Analysis

Hiker trail behavior data was summarized and statistical significance across study sites evaluated in the *gtsummary* package in R (version 4.5.1). P-values were obtained by the default statistical tests; Pearson’s Chi-squared test, Kruskal-Wallis rank sum test, and Fisher’s exact test were applied when appropriate. Chi-squared tests were used to assess whether categorical trail behaviors or conditions (i.e., time of day, walking through grass, bringing pets, and weather) impacted the likelihood of finding a tick during the tick check and t-tests were used for continuous behaviors (i.e., hike duration). To understand how engagement with tick stations may change over time, we visualized number of submissions per site per week and modeled weekly counts for each site using a generalized linear mixed model fit with *glmmTMB* (negative binomial, log link). We treated week as a categorical predictor to test for any difference in counts across weeks. The global effect of week was evaluated with a likelihood-ratio (χ^2^) test comparing the week model to a null model using the anova() base R function. To test whether the distribution of tick species and tick sex significantly differed by collection type, Fisher’s exact tests were calculated for each of the pairwise comparisons and p-values were adjusted by the Holm correction method to account for multiple comparisons. To understand how tick density estimates compared between surveillance by the tick station and active surveillance, we visualized ticks per survey submitted and ticks per 1000 m sampled by site and week. Since active surveillance was not conducted weekly, we further assessed the relationship between the two tick density estimates per half month and site and fit a linear model to the data using the lm() base R function.

We compared effort in terms of time spent obtaining a tick (effort per tick) using the tick station and active surveillance approaches. First, we used the fit linear model to predict ticks per survey submitted by tick stations with confidence intervals when tick densities by active surveillance were 1, 4, 16, and 64 ticks per 1000 meters sampled. For active sampling, we assumed that 1000 m would be sampled per visit and that it would take 150 minutes to sample 1000 m. For surveillance by the tick-check station, we assumed that tick stations would be visited twice per month and four hours per month would be spent on preparing supplies and entering survey data. We assumed a scenario where engagement with the station is low, defined as 15 surveys submitted per month, as well as a scenario with high engagement, defined as 30 surveys submitted per month. In addition, we proposed a hybrid scenario where a tick station is deployed, visited twice per month, and active surveillance is also conducted at each visit. We evaluated effort per tick for all scenarios when the one-way commute time to sites was 30 (near), 60 (moderate), and 120 minutes (far).

We grouped KAP survey participants into control or intervention groups by whether they read the informational signage. We calculated collinearity between survey questions using the Pearson correlation method (Supplementary Materials 4). For binomial correct vs. incorrect answers, odds ratios were used to quantify the association between reading the tick-check station sign and correctly answering the survey questions. Practices were categorized as either performed or not performed and evaluated between groups using odds ratios. For Likert scale questions, Fisher’s exact test was used to determine if distributional differences existed between the responses of the control and intervention groups. Participant responses for knowledge and practices were also assessed across demographic groups (i.e., residence history, age group). Questions relating to attitudes were not evaluated by demographic information. Correlations with residence history were assessed using odds ratios. Age groups were assessed using logistic regression, with overall model significance evaluated via a likelihood ratio chi-square test, followed by pairwise comparisons of age groups using estimated marginal means if significance was found. We also evaluated the relationship between perceived risk and behaviors using generalized linear mixed models followed by Wilcoxon rank sum tests.

### 2.9 Ethical Approval

The KAP survey, and its protocol, were exempted from review under category 2i by the Colorado State University International Review Board (IRB).

## 3 Results

### 3.1 Trail behavior and engagement

Overall, 262 surveys were submitted across the four tick-check stations between April and August 2024 (Table 2). The highest number of surveys were collected from the ELC (n = 113), and the fewest were collected from Mud Lake (n = 32). Surveys were designed to assess hiker trail behaviors expected to impact the likelihood of encountering a tick on trail. Trail behaviors were analyzed to determine the impact of: 1) time of day; 2) duration of hike; 3) walking through ankle length brush or grass; 4) recreating with a pet; and 5) weather conditions on the likelihood of finding a tick during the tick check.

**Table 2.**
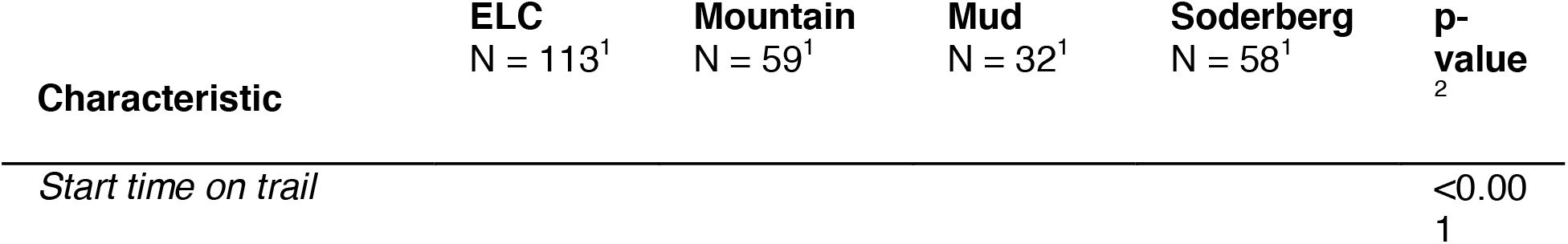
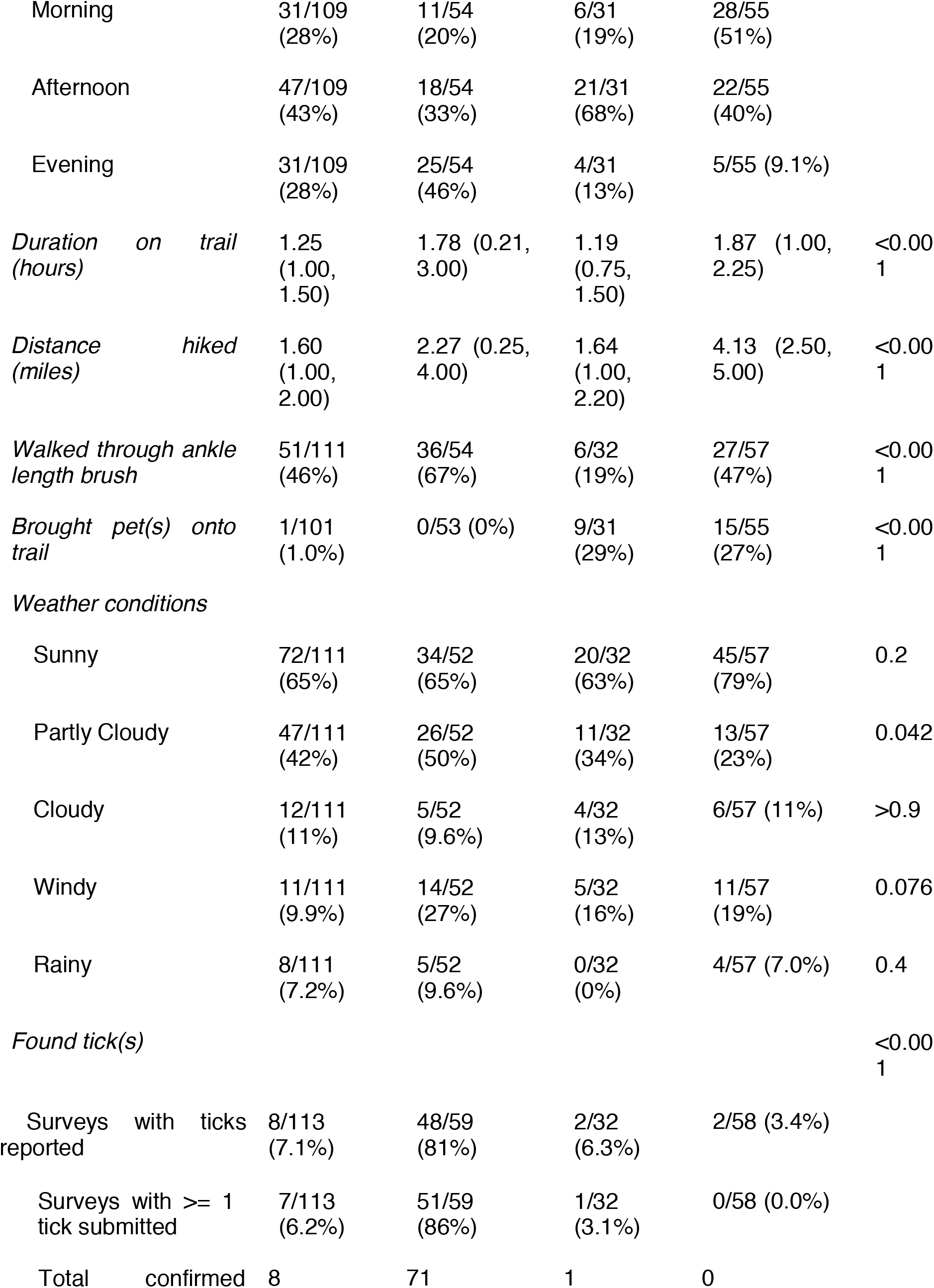
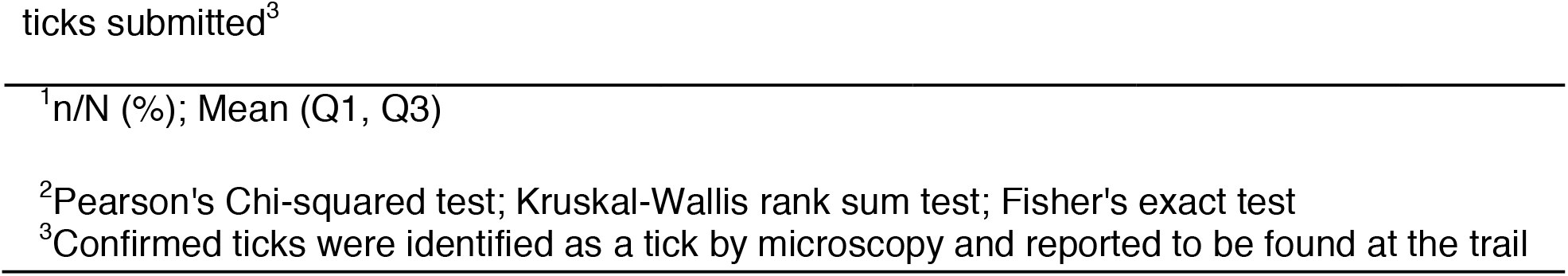
Hikers’ trail behavior data derived from survey submissions at the four tick-check stations. Percentages are derived from the number of surveys in which each characteristic was selected divided by the number of total surveys in which this section was filled out.

Start times on trails varied significantly by site (p < 0.001, Fisher’s exact test), with the afternoon being the most common start time reported at the ELC (47/109, 43%) and Mud Lake (21/31, 68%). The evening was the most popular start time among respondents at the CSU Mountain Campus (25/54, 46%), while the morning was the most common start time for respondents at Soderberg (28/55, 51%). There was a significant relationship between time of day and likelihood of finding a tick (p = 0.001, Chi-squared test), with evening hikes being the most likely time to find a tick.

Hike duration varied significantly by site (p < 0.001, Kruskal-Wallis test), with the longest average duration at Soderberg (1.87 hours) and the shortest average duration at Mud Lake (1.19 hours). Similarly, there was significant variation (p < 0.001, Kruskal-Wallis test) across sites for distance hiked. Soderberg had the highest average distance hiked, at 4.13 miles, while the ELC had the lowest, at 1.60 miles. There was not a significant relationship between hike duration and likelihood of finding a tick (p = 0.922, t-test).

We also assessed how many respondents walked through ankle length grass/brush during their hike (41). We found that over half of respondents reported walking through ankle length brush at the CSU Mountain Campus (36/54, 67%) which was the highest proportion of the four sites. The smallest proportion occurred at Mud Lake where roughly (6/32, 19%) of respondents reported walking through ankle length grass. The variation between these sites was significant (p < 0.001, Chi-square test). Individuals who walked through grass/brush were significantly more likely to encounter a tick than individuals who did not (p = 0.034, Chi-squared test).

Companion animals can also serve as tick hosts and increase the likelihood of encountering a tick (42). The variation in the percentage of respondents who recreated with pets varied significantly by site (p < 0.001, Fisher’s exact test). Notably, there were no pets reported at the CSU Mountain Campus (0/53, 0%). The highest percentage of respondents recreating with pets occurred at Mud Lake (9/31, 29%) while the highest overall number of pets reported at any of the sites was at Soderberg (15/55, 27%). Dogs are not permitted at the ELC, although we did receive one report of a pet at this site (1/101, 1.0%). We did not test statistical significance because pets were not allowed at multiple sites.

Weather conditions were analyzed by tallying the conditions reported on each survey to get the total amount of trail outings under each weather condition. These tallies were then divided by the total number of responses in which the respondent filled out the weather condition portion of the survey. Across all 4 sites, “sunny” weather was the most reported and there was not a significant difference in the weather between the sites (p > 0.05, Fisher’s exact test). No weather variable was significantly correlated with finding a tick (p > 0.05, Chi-squared test).

Additionally, we recorded the number of surveys that included ≥1 tick or indicated that a tick had been found despite no specimen provided with the survey. The variation in number of respondents who found ticks at each site was significant (p < 0.001, Fisher’s exact test) with the most ticks submitted from the CSU Mountain Campus (n = 71) and the least submitted at Soderberg (n = 0). Ticks that were reportedly found at locations other than the respective trailheads were excluded from analysis, as were submissions of specimens that were not identified as ticks.

We assessed community engagement with the tick-check stations over time at each of the four sites (Figure 3). Engagement was determined by the number of survey submissions per week. The first tick-check station was placed at the ELC during the week of April 15, 2024, while the other three stations were set up during the week of May 20, 2024. All stations received submissions throughout the study duration. There was no significant difference in surveys submitted per week for the ELC, Soderberg, and Mud Lake stations (p = 0.126, p = 0.146, p = 0.234, respectively, likelihood ratio test). There was a significant decrease (p = 0.009, likelihood ratio test) in submissions per week at the Mountain Campus, consistent with decreasing student enrollment as the summer session progressed.

**Figure 3.**
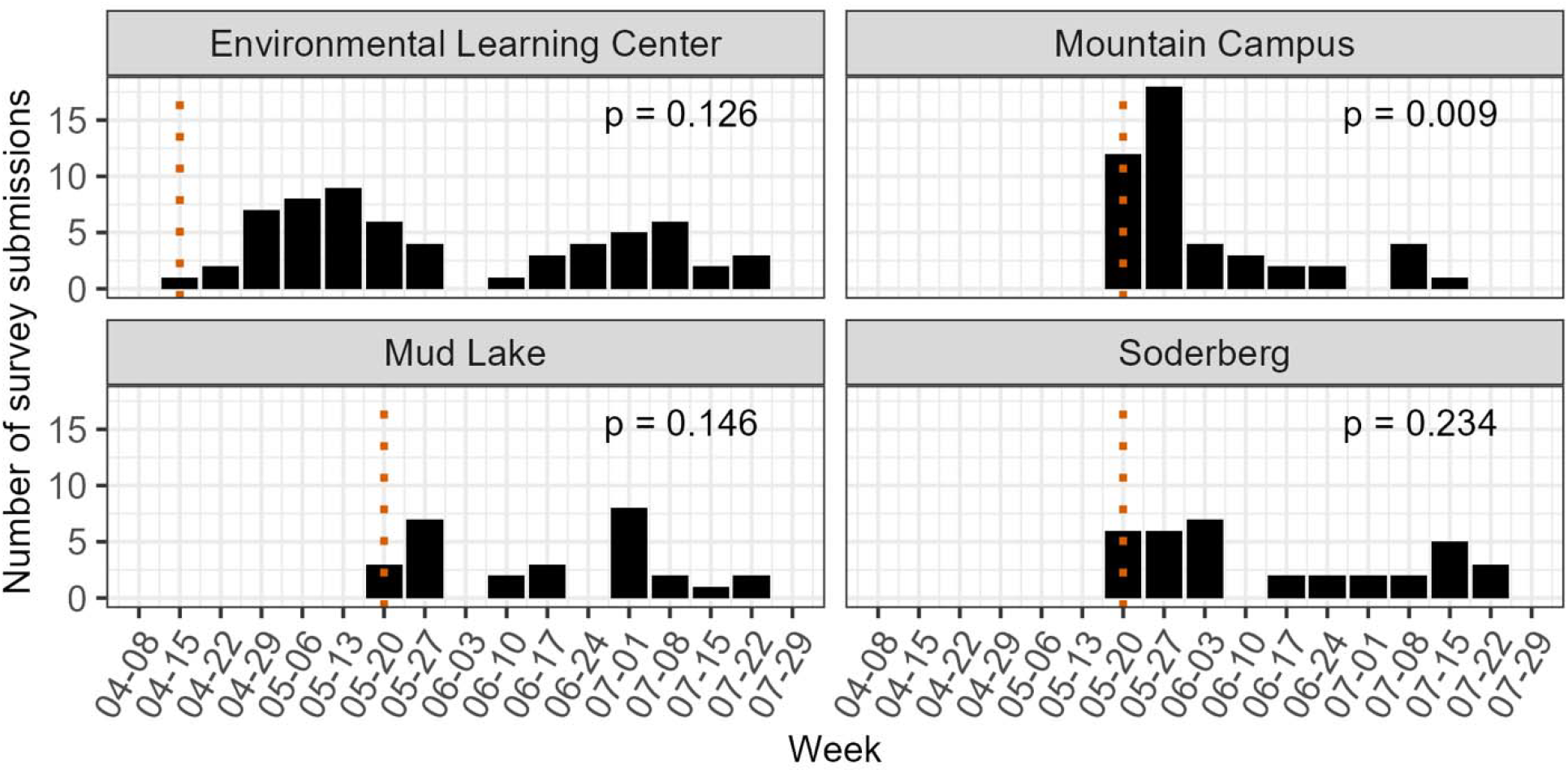
Engagement with tick-check stations over time. The dotted red line indicates the week that the station was installed, with regular check-ins occurring weekly or bi-weekly. Significance of weekly engagement across the study period was assessed using likelihood ratio tests.

### 3.2 Comparison of tick-check station and mail-in passive approaches

The CDPHE tick mail-in program is a statewide passive surveillance initiative that is divided amongst counties in Colorado (43). Since our study was restricted to Northern Colorado, we compared the results from our tick-check stations to mail-in data from Larimer County only. The raw CDPHE tick submission data for Larimer County is provided in the supplementary materials (Supplementary Materials 5). Although the CDPHE mail-in system was available year-round, and our tick-check stations were only available during the summer months, the tick submission date range was comparable between the two surveillance programs. From the date of the first submission to the date of the final submission for the mail-in system was 66 days and for our tick-check stations was 79 days. Therefore, even though our program was available to the community for a shorter period, the duration of engagement was comparable, indicating that the availability of the tick-check stations coincided with the period of relevant tick encounter risk. Further, the mail-in system captured more geographic variation compared to the tick-check stations (Supplementary Materials 6). Since our tick-check stations were only available at a limited number of specific sites, we were only able to capture a small amount of geographic variation.

Overall, there was a higher number of ticks submitted to our tick-check stations than through the mail-in program. We received 82 ticks from our tick-check stations, while 26 ticks were submitted to the CDPHE mail-in system (Supplementary Materials 7). However, there was a more even distribution of tick species (*D. andersoni*: *D. variabilis*) submitted through the mail-in system than the tick-check stations. The variability in tick species collected through the mail-in and the tick-check station was significant (p < 0.001, Fisher’s exact test) with more *D. variabilis* ticks submitted through the mail-in system (Supplementary Materials 7). In contrast, there was no significant difference in the distribution of life stages collected between the two methods, with adult females comprising the predominant stage for both (p = 0.395, Fisher’s exact test). No nymphs or larvae were collected by either surveillance method (Supplementary Materials 7).

### 3.3 Comparison of tick-check station and active surveillance

There was not a significant difference between tick species collected through active surveillance and tick-check station (p = 1, Fisher’s exact test) (Figure 4). Similarly, there was not a significant difference in tick life stage collected across the surveillance methods (p = 1, Fisher’s exact test). The predominant life stage collected by both surveillance methods was adult females. While a low number of nymphs were collected by active surveillance (n = 1), no nymphs were collected by either of the passive approaches and this difference was not significant (p > 0.1, Fisher’s exact test). No larvae were collected by either surveillance method.

**Figure 4.**
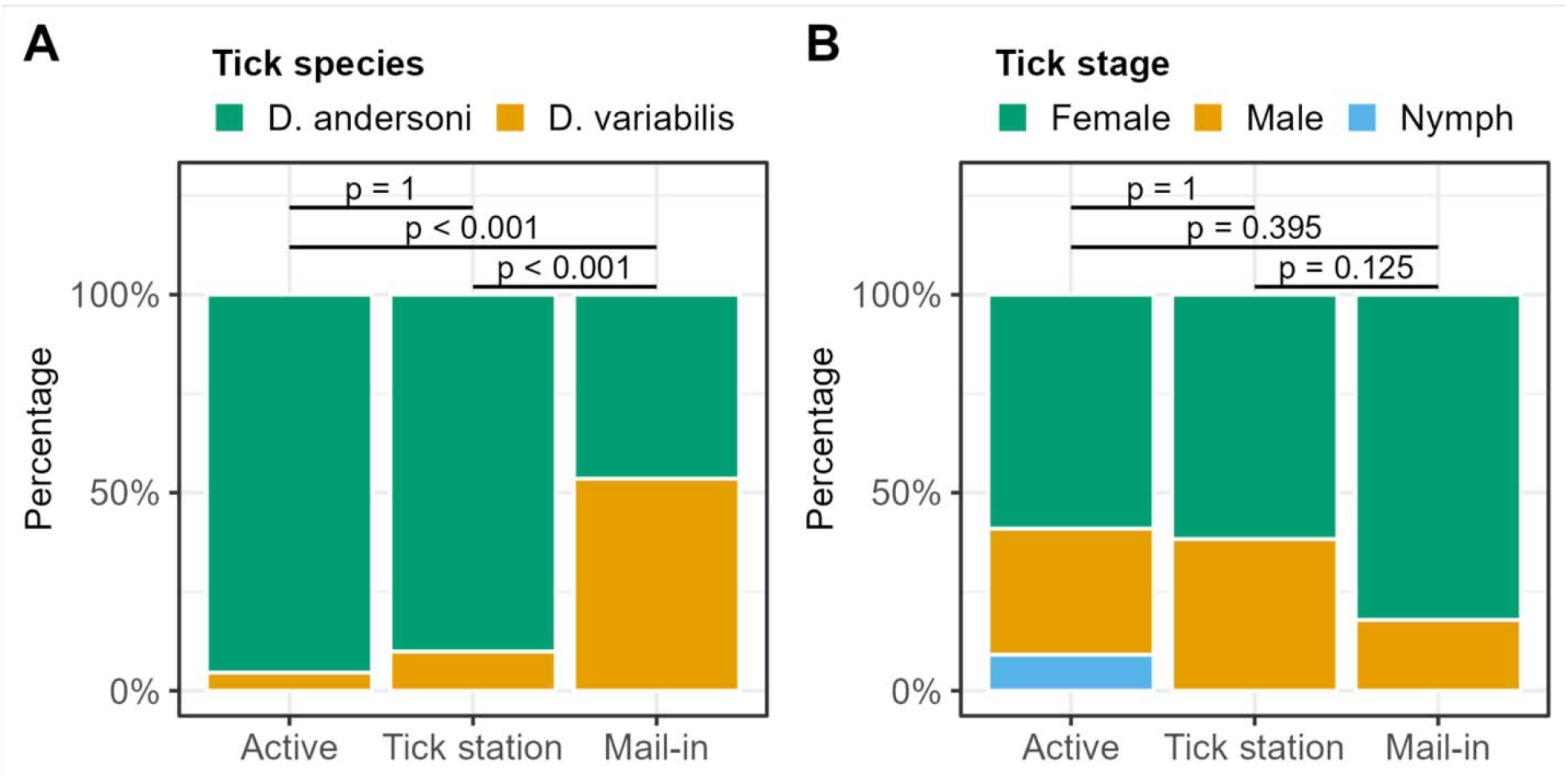
Tick species (A) and life stage (B) by surveillance approach. Significance was assessed using Fisher’s exact test.

We next compared tick density over time by active surveillance and tick-check station at the three sites where ticks were collected (Figure 5). No ticks were collected from either surveillance method at the Soderberg site, so it was excluded from further analysis. For our tick-check stations, density was calculated as the number of ticks collected per survey submitted. For active surveillance, density was calculated as the number of ticks collected per 1000 meters sampled. There was a distinct peak in tick density at the Mountain Campus site during the week of May 27^th^ for active surveillance, followed by a peak shortly thereafter in tick-check station density during the week of June 3^rd^. Notably, there was no active surveillance conducted at the Mountain Campus during the weeks of June 3^rd^ or June 10^th^. Trends in tick density were not evaluated at the ELC and Mud Lake due to the low number of ticks collected by either method at these sites.

**Figure 5.**
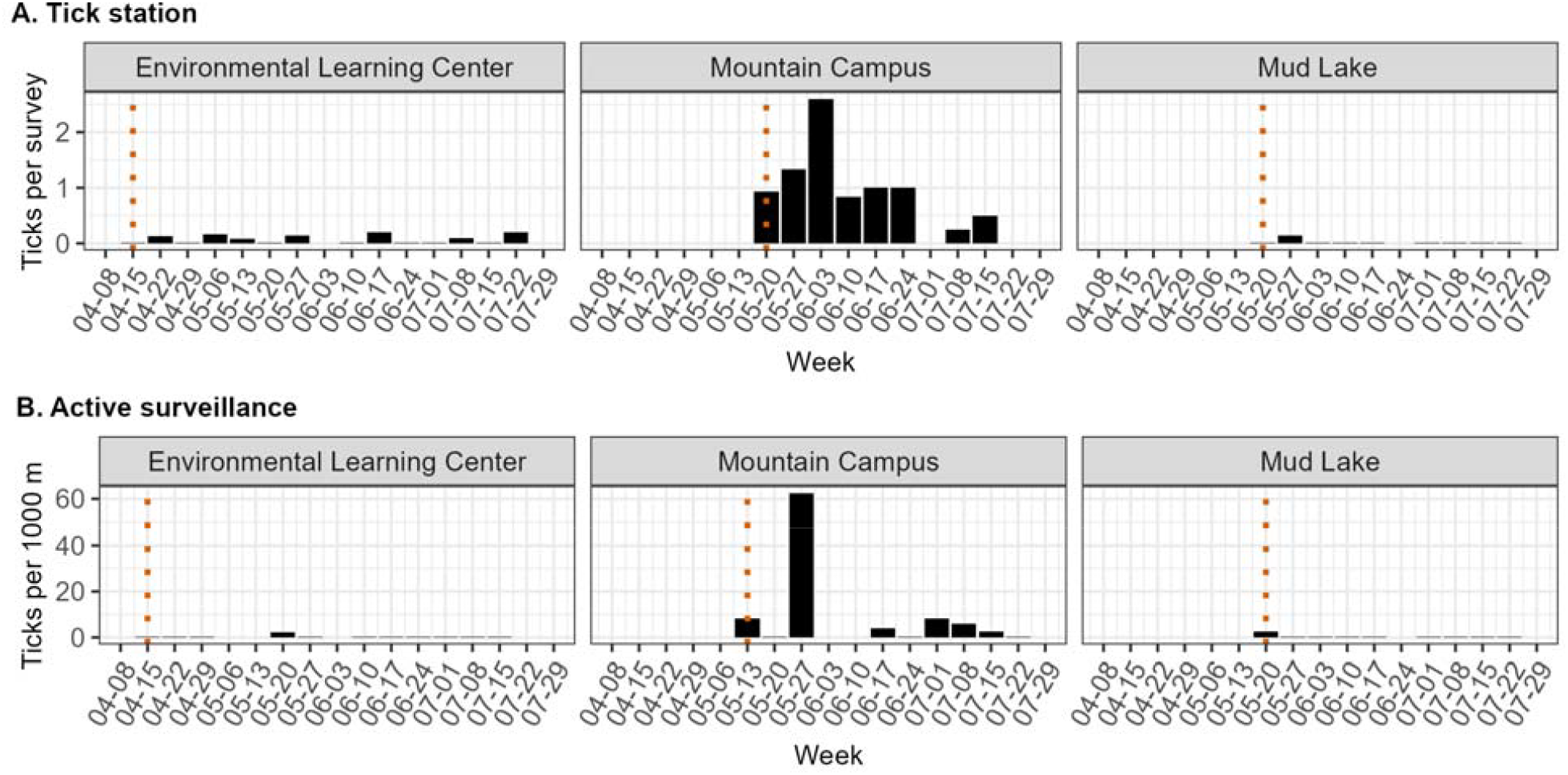
Observed tick density over time by surveillance type. (A) Tick check-station: Density measured as ticks per survey submitted. The dotted line indicates when the tick-check station was placed on-site. (B) Active surveillance: Density measured as ticks per 1000 m sampled. The dotted line indicates when active sampling began.

Next, we examined the relationship between tick densities derived from active surveillance and tick-check stations (Figure 6). There was a positive linear relationship between tick density estimated by active surveillance (ticks per 1000m) and tick density by tick-check station (ticks per survey submitted). Ticks per survey submitted increased by 0.13 for every 1-unit increase in ticks per 1000m (β = 0.13). This model explained 54.7% of the variance in ticks per survey submitted (R^2^ = 0.547).

**Figure 6.**
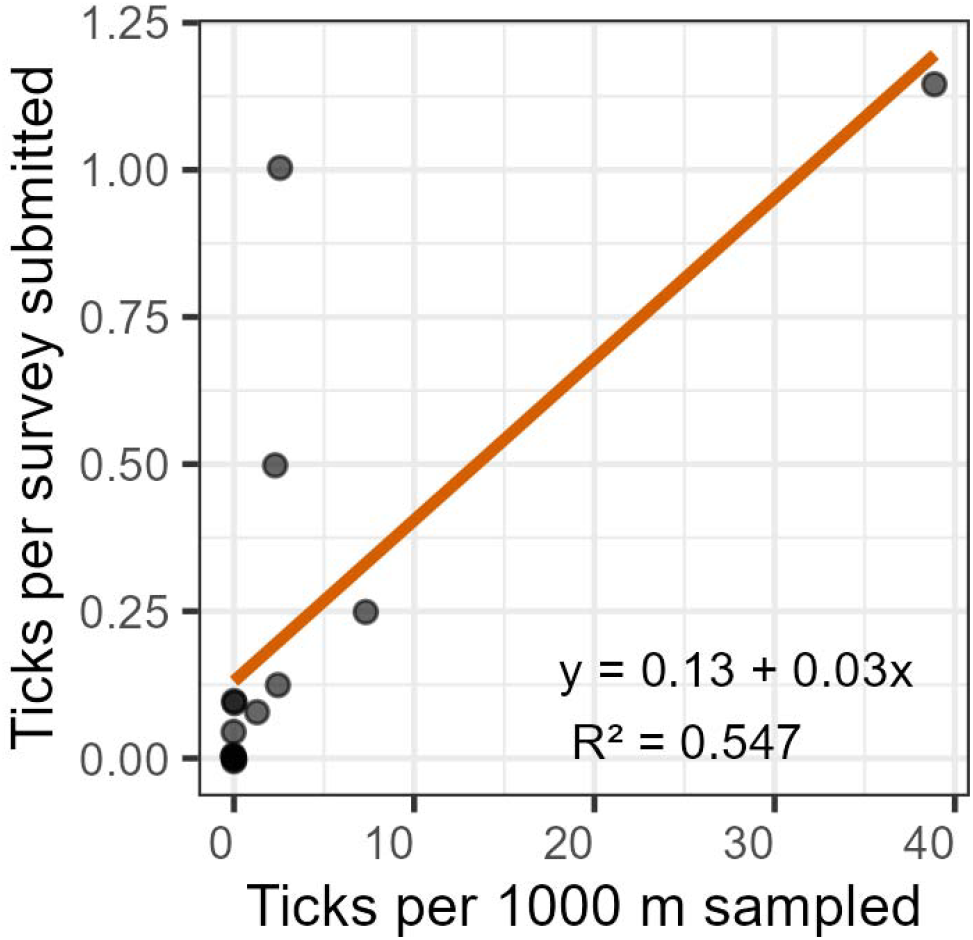
Comparison of tick densities measured by active surveillance (ticks per 1000 m sampled) and tick-check stations (ticks per survey submitted). Each point represents a specific site (ELC, Mountain Campus, or Mud Lake) over a half month duration. All points have the same level of transparency, and darker colors represent overlapping points. The orange line represents a linear regression.

### 3.4 Tick collection effort by surveillance approach

We compared relative effort (estimated in minutes) to collect one tick by each surveillance approach (Figure 7). Sampling methods were evaluated under simulated scenarios that varied by tick density (ticks per 1,000 m^2^), one-way commute time to the site (minutes), and collection type (tick-check station, active surveillance, or a hybrid approach). The hybrid approach was defined as a combination of tick-check station and active surveillance methods. The tick-check station and hybrid approaches were further stratified by community engagement level, with scenarios representing low engagement (15 surveys submitted per month) and high engagement (30 surveys submitted per month). Effort per tick was estimated in minutes, demonstrating the relative time investment needed to collect one tick by the given collection type and scenario. Several assumptions were made for these simulations. All activities were assumed to be performed by one person, and each site was visited twice per month. For tick-check stations, we assumed that four hours per month would be spent preparing surveys and recording results. For active surveillance, we assumed that 1,000 m^2^ were sampled per site visit and that each visit required approximately 2.5 hours.

**Figure 7.**
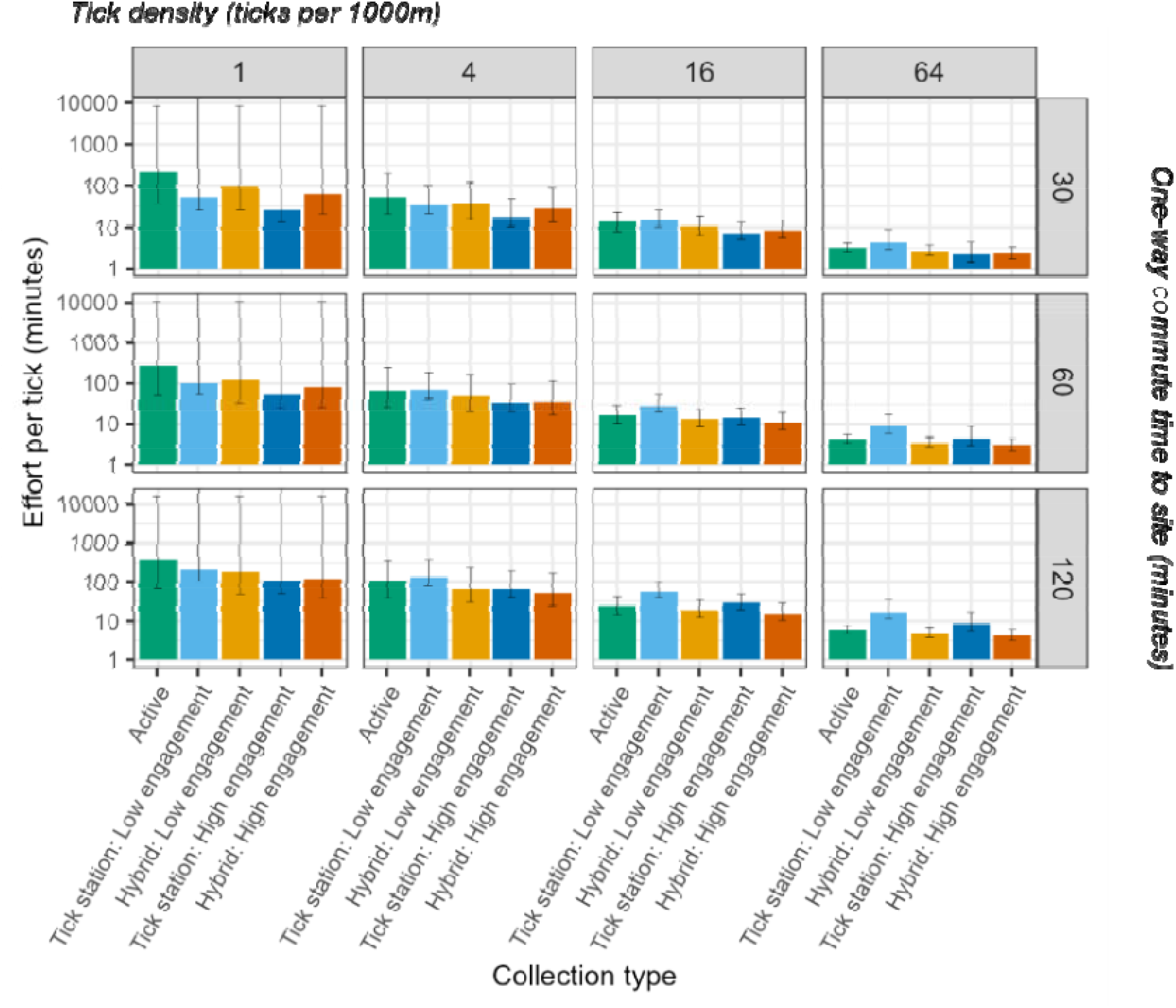
Simulated data examining the effects of site distance, sampling method, and tick density on the “effort per tick” measured in minutes. Error bars represent 95% confidence intervals.

Patterns of time efficiency differed across engagement levels, commute times, and tick densities (Figure 7). Under high engagement, tick-check stations alone consistently required the least effort when sampling sites were nearby (30-minute one-way commute) or when tick densities were low (1 tick per 1,000 m^2^). Under low engagement, tick-check stations required the least effort only under more limited conditions, specifically at tick densities of 1–4 ticks per 1,000 m^2^ with a 30-minute commute, or at 1 tick per 1,000 m^2^ with a 60-minute commute. In contrast, when tick densities were higher (≥4 ticks per 1,000 m^2^) and/or when commute times were longer (≥60 minutes), the hybrid approach was generally the most time-efficient strategy, regardless of engagement level. Active surveillance alone was not the most efficient approach under any scenario evaluated.

### 3.5 KAP Survey Results

We conducted 99 verbal KAP surveys from late June to early September 2024 (Table 3). Participants that were surveyed at sites without a tick-check station, or that reported not reading the signage posted at the tick-check station, were included in the control group (n = 69). Participants who reported reading the information at the tick-check station were included in the intervention group (n = 20). Participant demographic information is provided in the supplement (Supplementary Materials 3).

**Table 3.**
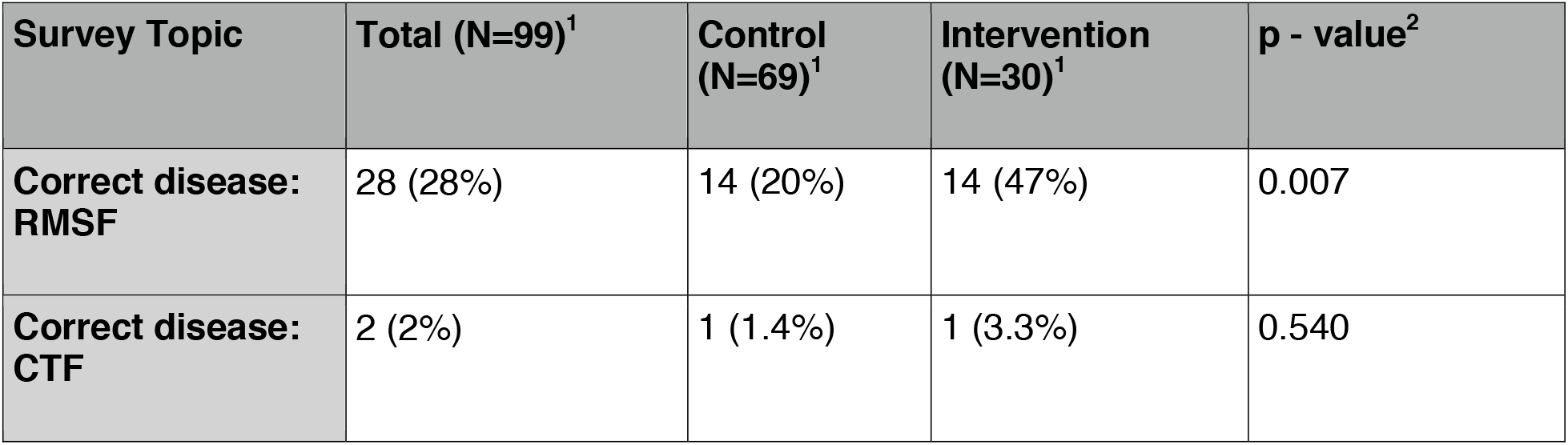
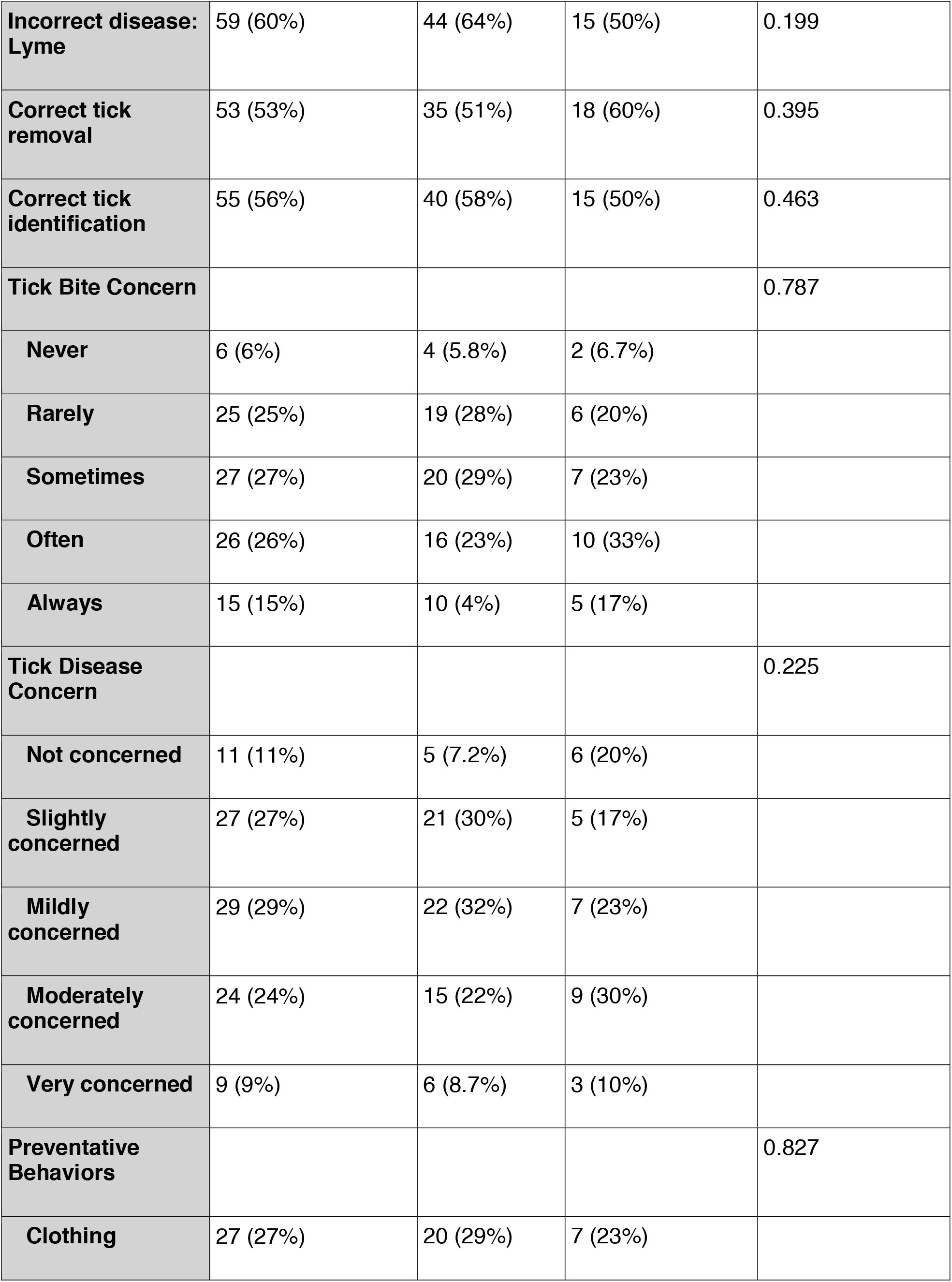
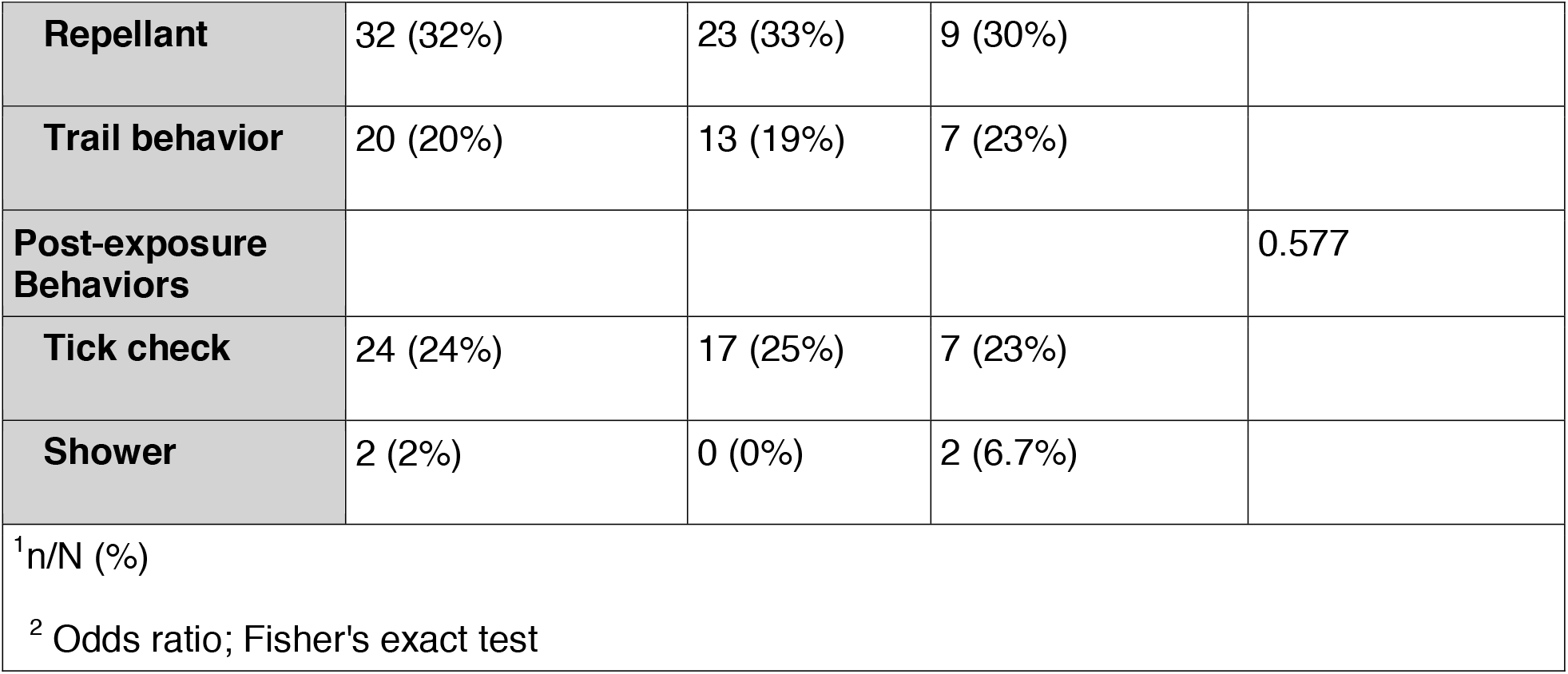
KAP survey results.

| Survey Topic | Total (N=99)<sup>1</sup> | Control (N=69)<sup>1</sup> | Intervention (N=30)<sup>1</sup> | p - value<sup>2</sup> |
| --- | --- | --- | --- | --- |
| **Correct disease: RMSF** | 28 (28%) | 14 (20%) | 14 (47%) | 0.007 |
| **Correct disease: CTF** | 2 (2%) | 1 (1.4%) | 1 (3.3%) | 0.540 |
| **Incorrect disease: Lyme** | 59 (60%) | 44 (64%) | 15 (50%) | 0.199 |
| **Correct tick removal** | 53 (53%) | 35 (51%) | 18 (60%) | 0.395 |
| **Correct tick identification** | 55 (56%) | 40 (58%) | 15 (50%) | 0.463 |
| **Tick Bite Concern** |  |  |  | 0.787 |
| **Never** | 6 (6%) | 4 (5.8%) | 2 (6.7%) |  |
| **Rarely** | 25 (25%) | 19 (28%) | 6 (20%) |  |
| **Sometimes** | 27 (27%) | 20 (29%) | 7 (23%) |  |
| **Often** | 26 (26%) | 16 (23%) | 10 (33%) |  |
| **Always** | 15 (15%) | 10 (4%) | 5 (17%) |  |
| **Tick Disease Concern** |  |  |  | 0.225 |
| **Not concerned** | 11 (11%) | 5 (7.2%) | 6 (20%) |  |
| **Slightly concerned** | 27 (27%) | 21 (30%) | 5 (17%) |  |
| **Mildly concerned** | 29 (29%) | 22 (32%) | 7 (23%) |  |
| **Moderately concerned** | 24 (24%) | 15 (22%) | 9 (30%) |  |
| **Very concerned** | 9 (9%) | 6 (8.7%) | 3 (10%) |  |
| **Preventative Behaviors** |  |  |  | 0.827 |
| **Clothing** | 27 (27%) | 20 (29%) | 7 (23%) |  |
| **Repellant** | 32 (32%) | 23 (33%) | 9 (30%) |  |
| **Trail behavior** | 20 (20%) | 13 (19%) | 7 (23%) |  |
| **Post-exposure Behaviors** |  |  |  | 0.577 |
| **Tick check** | 24 (24%) | 17 (25%) | 7 (23%) |  |
| **Shower** | 2 (2%) | 0 (0%) | 2 (6.7%) |  |
| <sup>1</sup>n/N (%) |  |  |  |  |
| <sup>2</sup> Odds ratio; Fisher's exact test |  |  |  |  |

#### Knowledge

Participants in the intervention group were significantly more likely to correctly identify RMSF as a TBD in Colorado than those in the control group (p = 0.007, Odds ratio). In the control group, 20% (14/69) of individuals correctly identified RMSF, compared with 47% (14/30) in the intervention group. Only two participants across both groups correctly identified CTF as a TBD in Colorado (control n = 1; intervention n = 1). More than half of participants in both groups incorrectly identified Lyme disease as a TBD in Colorado (59/99). While incorrect identification was lower in the intervention group (15/30, 50%) than in the control group (44/69, 64%), this difference was not statistically significant (p = 0.199, Odds ratio). Although a higher proportion of participants in the intervention group correctly identified an appropriate method for tick removal (18/30, 60%) compared with the control group (35/69, 51%), this difference was not statistically significant (p = 0.395, Odds ratio). There was also no significant difference in correct tick identification between groups (p = 0.463, Odds ratio), but a slightly higher proportion of participants in the control group correctly identified the ticks from images (40/69, 58%) compared with those in the intervention group (15/30, 50%).

#### Attitudes

Overall, most participants reported *rarely, sometimes*, or *often* having concerns about tick bites, with fewer respondents selecting *never* or *always*. This distribution was consistent across both groups, and no significant differences were observed between groups (p = 0.787, Fisher’s exact test). For concerns about TBD, overall responses were likewise concentrated in the mid-range categories (*slightly, mildly*, or *moderately concerned)*, with fewer respondents selecting *not concerned* or *very concerned*.

#### Practices

Practices related to tick bite and TBD exposure prevention were categorized as preventative behaviors (e.g., wearing protective clothing, applying repellant, and remaining on designated trails) or post-exposure behaviors (e.g., conducting a tick check, showering after outdoor activity). Across both groups, the most commonly reported preventative behavior was applying repellent (32/99, 32%), followed by wearing appropriate clothing (27/99, 27%) and staying on the trail (20/99, 20%). The most commonly reported post-exposure behavior was conducting a tick check (24/99, 24%). The proportions of participants reporting each preventive and post-exposure practice were similar between the control and intervention groups, and no statistically significant differences were observed for any individual behavior (p > 0.05, Odds ratio; Supplementary Materials 8).

Additionally, disease concern was significantly correlated with practicing any protective behavior across both groups (p = 0.03, Wilcoxon rank sum test). The proportion of individuals reporting at least one protective behavior generally increased with higher levels of concern: 73% (8/11) among those reporting “not concerned”; 65% (17/26) “slightly concerned”, 69% (20/29) “mildly concerned”, 88% (21/24) “moderately concerned”, 100% (9/9) among those reporting “very concerned”.

Finally, to explore potential demographic covariates associated with survey responses, we compared participant demographic information with KAP outcomes (Supplementary Materials 8). We found that participants who were current or previous residents of a Midwestern state were more likely to report engaging in post-exposure prevention practices (15/40, 37.5%) than participants who have never resided in a Midwestern state (p = 0.036, Odds ratio). Additionally, we found that significantly more individuals aged 36-45 had knowledge of correct tick removal practices than those aged 46-55 (p = 0.022, likelihood ratio chi-squared). Lastly, we found that individuals aged 26-35 were more likely to misidentify Lyme disease as a Colorado TBD than those aged 55+ (p = 0.019, likelihood ratio chi-squared).

## 4 Discussion

With the geographical range expansion of ticks in the U.S. and rise in reported TBD cases, effective surveillance methods are now more important than ever (1,4,9–11). The use of combined active tick dragging and passive tick-check stations at our study sites allowed us to evaluate the effectiveness and effort required for each method individually as well as combined via a hybrid approach. To our knowledge, this is the first systematic evaluation of tick-check stations as tools for surveillance and public education. We found that estimates of tick densities, species composition, and life stage abundances derived from tick-check station data were reflective of estimates from concurrent active surveillance. Moreover, tick-check stations require less effort per tick than active surveillance in several scenarios, particularly in areas of low tick density and/or when study sites are nearby. In areas of high tick density and/or when study sites are far, a hybrid approach combining tick-check stations with active surveillance was the most time-efficient. In addition to their surveillance value, the informational signage at tick-check stations may improve TBD knowledge; the value of this benefit is underscored by the low baseline TBD knowledge observed in our study. Altogether, these results suggest that incorporating tick-check stations into surveillance programs can provide reliable estimates of tick encounter risk, improve time efficiency across a range of contexts, and enhance public health education.

Passive surveillance approaches are commonly integrated into state tick surveillance programs, but their effectiveness for estimating tick-borne disease risk remains unclear. Existing evaluations of passive surveillance approaches are sparsely available and uneven in scope. One study found that distributing tick collection kits at county-level hubs resulted in the collection of more ticks than active surveillance but did not compare tick density estimates between approaches (44). Another study comparing a state mail-in system with active surveillance found that passive submissions captured county-level patterns of tick and pathogen presence comparable to active surveillance; however, no relationship was observed between estimates of tick abundance or the abundance of infected ticks across approaches (14). In our study, tick-check stations collected more ticks within Larimer County than either concurrent active surveillance or the state mail-in system. Tick-check stations also provided geographically accurate estimates of tick populations, with species composition and life stage distributions comparable to those obtained through active sampling. Moreover, spatiotemporal tick densities estimated from tick-check stations were linearly correlated with density estimates from active surveillance, accounting for roughly half of the observed variation and potentially offering a more realistic measure of human exposure risk. Although the state’s mail-in system captured ticks across a broader geographical range, tick-check stations yielded higher tick counts, potentially increasing statistical power to detect pathogens circulating at low prevalence. Together, these findings suggest that tick-check stations represent a valuable complement to existing surveillance programs by providing spatiotemporal estimates of tick densities that scale with active surveillance and increasing sample sizes for estimating tick-borne disease risk.

Public health departments often have limited or inconsistent resources to sustain systematic tick surveillance, even as demand for tick distribution and pathogen risk information grows (5). As a result, many agencies rely on passive mail-in or voluntary tick submission systems as key components of tick surveillance activities because they are comparatively cost-effective (45). However, such submission programs are typically opportunistic rather than systematic, producing data that can be spatially uneven and difficult to compare across locations or over time. In this context, tick-check stations may offer a useful complement to existing surveillance programs due to their efficiency and greater standardization and geographic specificity. With modest upfront investment, stations can be constructed and subsequently maintained with relatively low recurring costs. Across many scenarios, tick-check stations yielded a higher efficiency than active surveillance, particularly when tick densities were low or field sites were far away. Moreover, implementing a hybrid model, where active dragging was conducted during routine station visits to collect submissions and replenish supplies, further reduced effort per tick across a range of scenarios. Importantly, because submissions were paired with survey data, stations also enabled assessment of encounter risk factors. We observed higher likelihood of tick encounters in the evening and among individuals who reported leaving designated trails, highlighting the value of this approach for identifying behavioral, environmental, and temporal drivers of exposure risk. Together, these findings suggest that integrating tick-check stations into existing surveillance frameworks may enhance efficiency, generate standardized estimates of tick encounter risk, and promote public engagement.

Personal protection is the primary intervention for preventing tick-borne diseases, underscoring the importance of understanding how public health campaigns lead to behavioral changes. In examining the knowledge component, awareness of non-Lyme TBDs was relatively low in our study, consistent with previous studies across the United States (24,46–48). More participants incorrectly identified Lyme disease as endemic to Colorado (59/99, 60%) than correctly identified RMSF (28/99, 28%) or CTF (2/99, 2%) as endemic to the state. The lack of awareness of CTF is particularly striking given that it is the most prevalent TBD in Colorado (49). Encouragingly, participants exposed to the tick-stations were more likely to correctly identify RMSF as a disease present in Colorado than those in the control group, suggesting that this intervention may improve disease specific awareness. Approximately half of respondents were able to correctly describe how to safely remove a tick and identify tick images, a level of knowledge that was lower than that reported in other studies (50,51). Collectively, these findings highlight opportunities for targeted educational interventions to address gaps in knowledge of locally relevant TBDs and suggest that tick-check stations represent one practical tool to help fill these gaps.

Within the KAP framework, attitudes encapsulates the ways which perceived risk, benefits, barriers, and personal experiences play a central role in determining whether knowledge translates into motivation for adopting protective behaviors (52–55). Consistent with this framework, we found that greater concern for TBD diseases was associated with a higher likelihood of engaging in protective behaviors. This finding aligns with previous studies in TBD contexts, which have reported that individuals with higher perceived prevalence, severity, or likelihood of contracting a TBD were more likely to engage in preventive behaviors (cite). At the same time, most participants in our study reported only modest concern about tick bites and TBD risk, a perception that may be proportionate to the relatively low incidence of TBDs in Colorado. This level of concern differs from that reported in higher incidence regions, particularly Lyme disease-endemic areas, where a greater perceived risk of contracting a TBD is documented (52,56). Altogether, these findings suggest that the lower perceived risk we observed in our study may be a factor in the relatively low adoption of protective behaviors reported.

Because personal protective behaviors represent the primary means of preventing TBD, understanding adoption of these practices is critical to reducing disease risk. Among them, performing a tick check is considered the gold standard for preventing tick attachment and TBD transmission (57–59); however, fewer than 25% of respondents reported performing tick checks after recreating outdoors. This value is much lower than from studies in Lyme disease endemic areas where tick-check practices often range from 45%-68% (56,60,61). Notably, participants with a history of residence in the Midwest were significantly more likely to perform tick checks than individuals who had never lived in the region, suggesting that sustained exposure to public health messaging in higher-incidence settings may influence long-term adoption of preventative behaviors. In contrast, more participants (n = 32/99, 32%) reported using repellents as a tick prevention strategy; this rate is comparable to that observed in Lyme-endemic areas (56,60,61). The divergence between relatively low tick-check adherence and more common repellent use suggests that motivation to engage in preventative practices exists, but awareness of tick checks as an effective strategy may be limited. Although this mechanism was not explicitly assessed in this study, this pattern highlights an additional opportunity for targeted outreach and education focused on promoting awareness of evidence-based prevention strategies. Although our study design did not allow us to determine whether exposure to our tick-check station translated into sustained shifts in attitudes or practices, we established baseline measures of risk perception and preventive behaviors and identified clear opportunities to strengthen disease-specific awareness and promote behaviors that reduce TBD risk.

When utilizing tick-check stations for passive surveillance, several considerations extend beyond time and cost. Our primary goal for developing passive tick-check stations was to characterize tick-human encounters at specific trails and geographic locations. However, some submissions were inconsistent with this aim and could not be used to characterize tick exposure among trail users. For example, some individuals submitted ticks collected outside the study trails, submitted non-tick arthropods, or reported finding a tick in the survey without submitting a specimen (and vice versa). Although excluded from the final analysis, these submissions nonetheless suggest that the tick-check stations foster community engagement and interest in tick surveillance. Second, return rates for tick collection kits are another consideration. By the end of the five-month study period, 41% of distributed kits were returned; however, these kits yielded a substantial number of ticks (n = 80). Future implementation of similar passive collection methods should account for potential loss of supplies over time, associated replenishment costs, and sustainability challenges. Third, placement of tick-check stations should align with study objectives. If the goal is to characterize exposure across a broader landscape, stations should be placed in areas representative of where human–tick encounters are most likely to occur. Notably, the species composition of ticks submitted to CDPHE from Larimer County differed from that observed at our tick-check stations, suggesting that station placement may not have fully captured broader county-level exposure patterns. In addition, stations placed at high-traffic trailheads may maximize participation and specimen return, though they may not provide regionally representative estimates of tick community composition. Thoughtful selection of station locations is therefore essential to balance representativeness, engagement, and surveillance yield.

This study had certain limitations. As a pilot study, surveillance was geographically constrained, with data collected from a limited number of trails in Northern Colorado. Community engagement levels strongly influence the effectiveness of passive surveillance tools, and community participation may vary across states or regions. Outdoor recreation plays a large role in community leisure activities and culture in Northern Colorado, and the recreational spaces we selected for this study were located in close proximity to multiple university campuses (62). These contextual factors may have positively influenced the level of engagement our tick-check stations received, and thus their effectiveness. Additionally, Colorado is considered to be a low tick-incidence state (63,64). As a result, our findings may not be directly generalizable to higher incidence regions where risk perception, exposure frequency, and susceptibility to messaging fatigue or desensitization may differ. In addition, we used time as a proxy to measure effort per tick when evaluating the effectiveness of our tick-check stations under different scenarios. We did not perform a cost analysis as fluctuating prices for materials and hourly employee wages differ over time and across regions. Finally, individuals who found a tick during their tick-check may be more likely to submit a survey, limiting our ability to define a true denominator for estimating encounter rates. Future studies could address this limitation by applying Bayesian approaches to estimate the underlying denominator or by integrating trail-use metrics, such as motion sensors or traffic counters, to better approximate encounter rates. Despite these limitations, this pilot study provides important preliminary insights into the feasibility and utility of passive tick-check stations as tools for surveillance and public education.

This study compared the effectiveness of tick-check stations to traditional mail-in and active surveillance approaches. Our findings suggest that tick-check stations can engage community members in ways that enhance surveillance while simultaneously increasing TBD awareness. By leveraging community participation, this model may reduce the personnel effort typically required for active tick surveillance while generating high geographic resolution data, improving understanding of tick-human encounter risk factors, and providing community education opportunities regarding TBD risk and prevention. Overall, we conclude that tick-check stations are a promising tool that can be integrated into public health organizations’ tick surveillance programs.

## Acknowledgements

We thank Foram Raval, Sofia Christensen, Logan Lowe, Jon Wegryn, Jake Brisnehan, and Brooke Shenkenberg for their assistance with tick surveillance and in-person KAP surveys. We gratefully acknowledge Boulder County Parks & Open Space, the City of Fort Collins, Larimer County Department of Natural Resources, the CSU Mountain Campus, and the CSU Environmental Learning Center for providing permission, site access, and logistical support for this research. This work was supported by the Colorado State University One Health Institute, the Infectious Disease Research and Response Training Program (NIH NRSA T32AI162691), and the Rockies and High Plains Vector-borne Diseases Center. The Centers for Disease Control and Prevention, Department of Health and Human Services, provided financial support for this project. The award provided 20% of total costs (US$5,675,000.00). The contents are those of the authors. They may not reflect the policies of the Department of Health and Human Services or the US government.

## Author contribution statement

EKH, KR and EHS conceived and designed the study, supervised the project, and secured funding. LD constructed the tick-check stations. CF, LD, and SG maintained the tick-check stations and conducted active tick surveillance and in-person surveys. NK and SC designed the KAP surveys. CMR led the mail-in tick submission program. EHS and SG performed data analysis. CF, LD, SG, and EHS drafted the initial manuscript. All authors contributed to manuscript revision and approved the final version.

## Notes

**Funding information:** This work was supported by the Colorado State University One Health Institute, the Infectious Disease Research and Response Training Program (NIH NRSA T32AI162691), and the Rockies and High Plains Vector-borne Diseases Center. The Centers for Disease Control and Prevention, Department of Health and Human Services, provided financial support for this project. The award provided 20% of total costs (US$5,675,000.00). The contents are those of the authors. They may not reflect the policies of the Department of Health and Human Services or the US government.

### Competing Interest Statement

The authors have declared no competing interest.

