## Supplementary material 1 for "Comparison of tick surveillance approaches: Utilizing trailhead tick-check stations to support tick surveillance and community education"

| **Material** | **Quantity** | **Price*** | **Total Cost** | **Purpose** | **Ordering Information** |
| --- | --- | --- | --- | --- | --- |
| Pressure treated pine lumber 2" x 4" x 8' | 3 | $6.18 | $18.54 | Used to construct the frame | Home Depot (SKU #1001802203) |
| Pressure treated pine lumber 2" x 2" x 8' | 2 | $5.38 | $10.76 | Used to attach poster | Home Depot (SKU #384090) |
| 19-gallon heavy duty rope handle plastic tub | 1 | $28.50 | $28.50 | Holds the main post and all attachments, makes unit easier to transport | Home Depot (SKU #1011131615) |
| 50lb bags of fast setting concrete mix | 2 | $6.91 | $13.82 | Keeps everything in place within 19-gallon plastic tub | Home Depot (SKU #842303) |
| 0.5 cubic feet landscape rock | 1 | $7.97 | $7.97 | Weighs down unit once at intended site, aesthetics | Home Depot (SKU #440943) |
| 3.5" exterior flat head deck wood screws contruction torx T-Star Plus | 1 | $12.97 | $12.97 | Used to construct the frame and add attachments | Home Depot (SKU #1006213454) |
| 2.5" exterior flat head deck wood screws contruction torx T-Star Plus | 1 | $12.97 | $12.97 | Used to construct the frame and add attachments | Home Depot (SKU #1006213445) |
| 1.25" exterior flat head deck wood screws contruction torx T-Star Plus | 1 | $12.97 | $12.97 | Used to construct the frame and add attachments | Home Depot (SKU #1004031051) |
| Tan Sheffield Field Box 11.5" x 5.06" x 7.25" | 1 | $16.89 | $16.89 | Stores tick removal kits/surveys | Amazon (ASIN #B07B1WRBG8) |
| Weatherproof Wall Mount Locking Mailbox | 1 | $32.43 | $32.43 | Stores trail surveys | Amazon (ASIN #B01LZH7BTC) |
| Outdoor Brochure Holder | 2 | $16.99 | $33.98 | Stores completed surveys/any tick samples | Amazon (ASIN #B01H10JUL6) |

*Note price is from Summer 2024 in Colorado and is subject to change

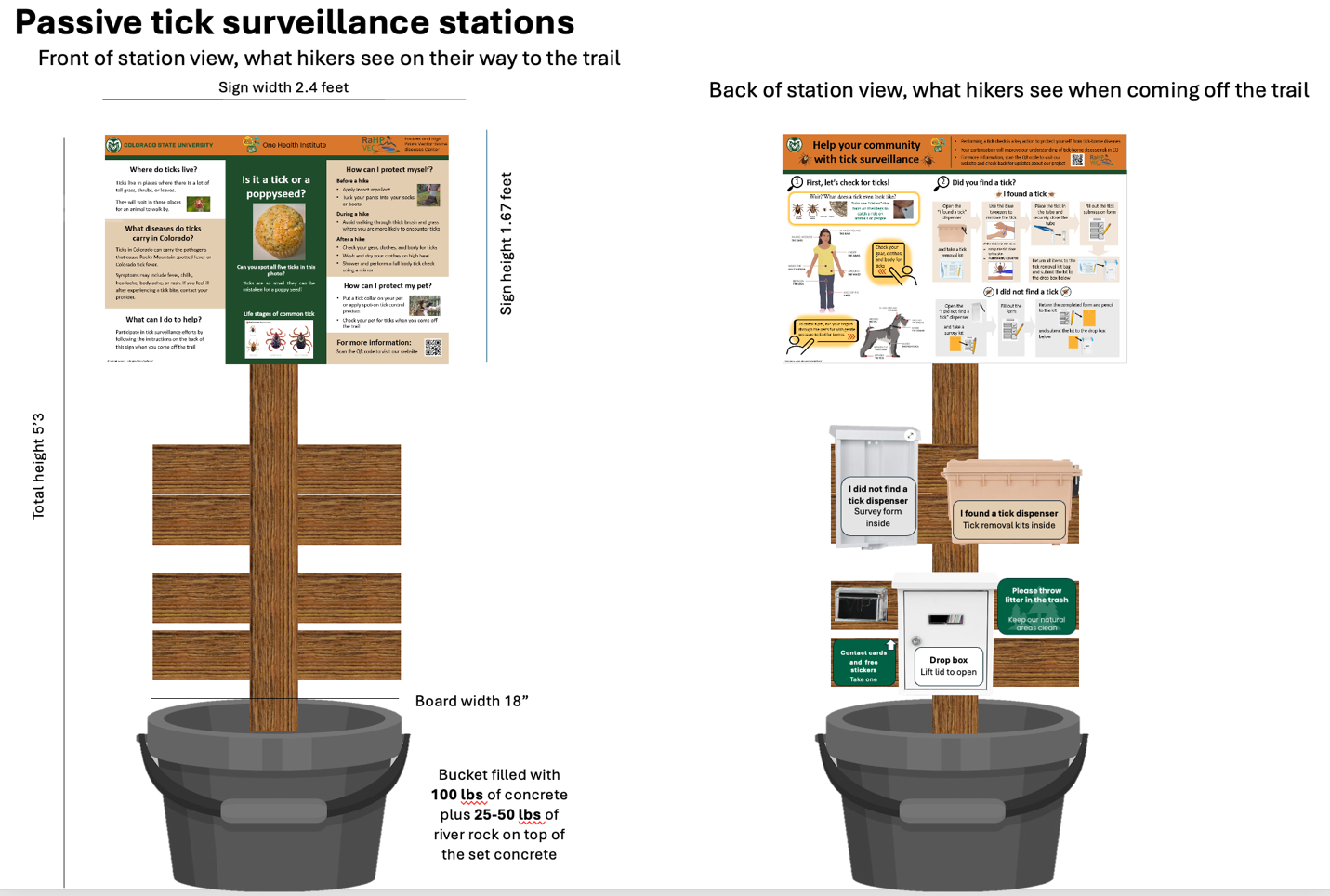

**Protocol for construction of passive surveillance tick collection stations:**

- **Constructing the frame (*please note exact positions on main post are subject to change based on size and shape of poster, mounted lockbox, and survey/tick collection kit dispensers)**
  - Place two pressure treated 2” x 4” x 8’ lumber pieces on a hard surface
  - Cut into eight pieces 2” x 4” x 2’
  - Place second 2” x 4” x 8’ on hard surface to serve as **Main Post**
  - Place one of the cut pieces approximately 45” from the bottom of main post and drill two 2.5” wood screws through both pieces
  - Place another of the cut pieces approximately 2.5” up the top of the other cut piece and drill two 2.5” wood screws through both the cut piece and the main post
  - Place third cut piece approximately 8” up from second cut piece on main post and drill two 2.5” wood screws through both the cut piece and the main post
  - Place fourth cut piece directly above third cut piece and drill to main post with two 2.5” wood screws through both the cut piece and the main post

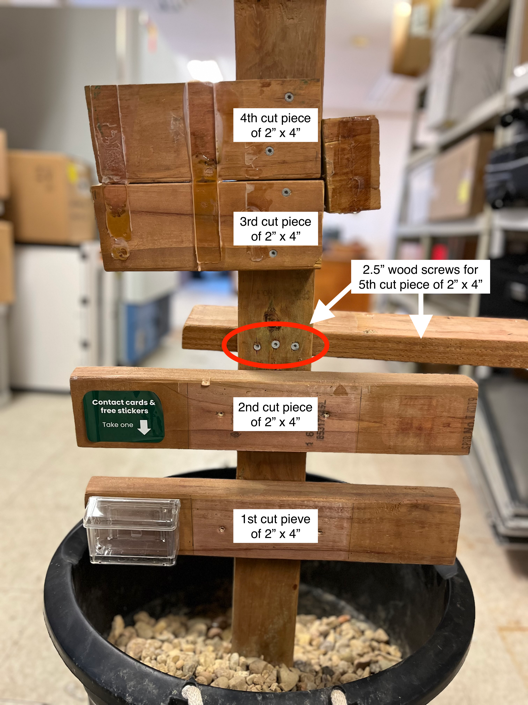

- Flip main post and attached pieces over
- Attach fifth cut piece to flipped main post approximately 40” from the bottom of the main post using three 2.5” wood screws that are screwed in from other side of the main post into the 2” side of the piece (ensuring the 4” wide side is facing up and can have Sheffield Field Box attached to it.

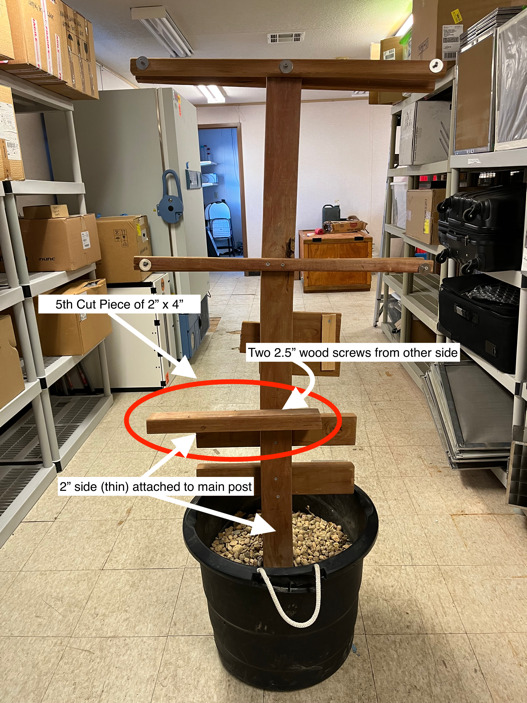

- Store other three cut pieces as replacements
- Place two 2” x 2” x 8’ lumber pieces on hard surface and cut into four pieces 2” x 2” x 3’
- Place one of these cut pieces on either side on the top of the main post and drill two 2.5” wood screws through both pieces
- Place remaining two pieces approximately 19-20” from top of main post and drill one on each side using two 2.5” wood screws for each

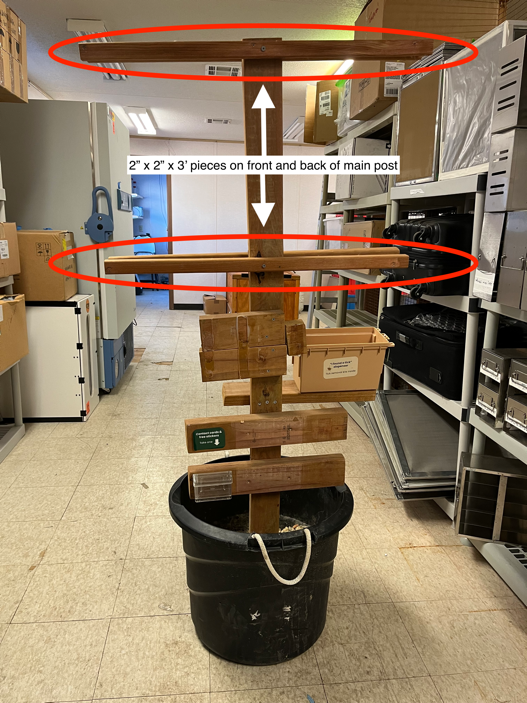

- **Mounting lockbox, survey/tick collection kit dispensers, and poster**
  - Drill four 1.25” wood screws into lower 2” x 4” pieces on main post (two screws on top piece and two screws on the bottom piece) spaced to fit the holes in the back of the lockbox
  - Carefully mount lockbox onto screws and make sure it is securely attached
  - Using wood glue, attach survey dispenser to the 2” x 4” pieces above the lockbox (adjust to ensure dispenser is centered)
  - Flip main post over to back side with one wood piece attached
  - Drill four 1.25” wood screws through the bottom of the Sheffield Field Box (to serve as tick collection kit dispenser) and into the top of the fifth cut 2” x 4”

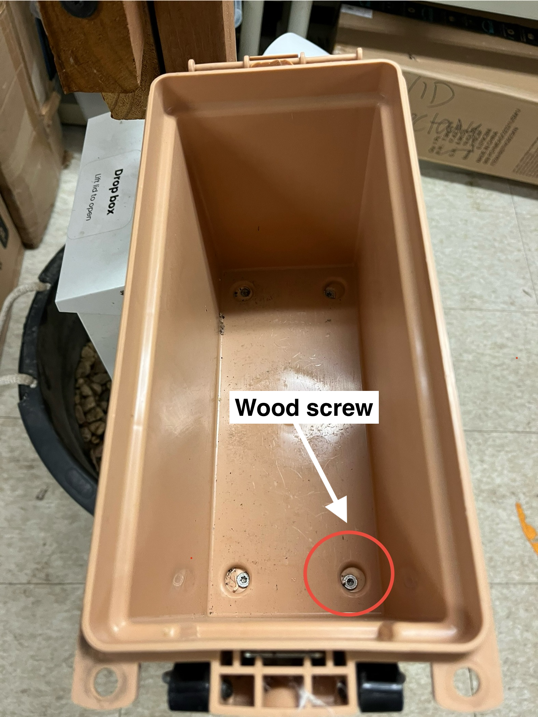

- Line up poster and attach to 2” x 2” pieces at top of main post at each of the corners using eight 1.25” wood screws
- **Concreting Station into plastic tub**
  - Place main post centered in the middle of the plastic tub
  - Pour in one 50 lb. bag of fast setting concrete mix
  - Add water until concrete mix is saturated and mix
  - Pour in second 50 lb. bag of fast setting concrete mix
  - Add water until concrete mix is saturated and mix
  - Hold main post in place until concrete has dried enough to keep it in place (5-10 minutes).
  - Add 25-50lbs of landscape rock on top of concrete (add when station is on-site as added weight makes transport difficult).

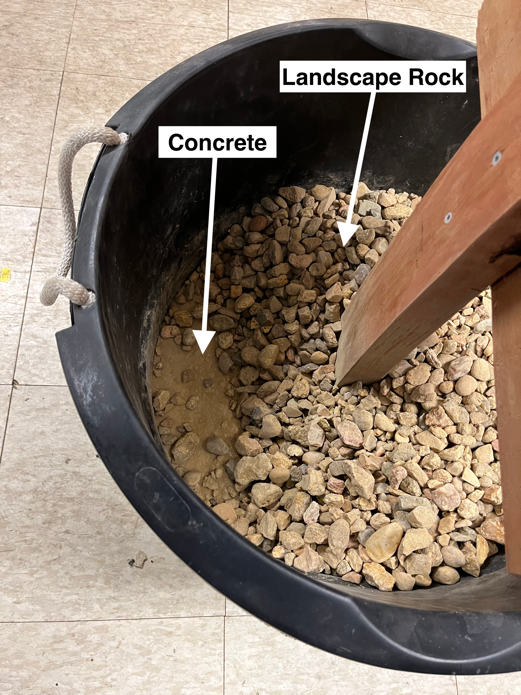
