## Supplementary material 2 for "Comparison of tick surveillance approaches: Utilizing trailhead tick-check stations to support tick surveillance and community education"

**Survey Consent Script:**
In conversational style, …

Hello, my name is (*investigator's name*) and I am a (*student researcher* or *researcher*) from Colorado State University. We are conducting a study on the distribution of ticks, and individual's knowledge about tick-borne diseases. We have a brief survey that will take approximately 5 minutes. There are no known risks or direct benefits to you, but we hope to better understand community knowledge about ticks through this survey.

If you decide to participate in our study, you may withdraw your consent and stop participation at any time without penalty. We will not collect your name or personal identifiers.

Would you like to participate?

If yes: Proceed.
If no: Thank you for your time.

In the form of a contact card, we will provide participants with the PI's names and contact information, the Participant's Rights contact information (; 970-491-1553.), funding information, and the URL link and QR code that directs to the project website (https://col.st/2rMO9) where participants can find more information about the project.

**Survey Questions:**
*Explanatory variables*

1. Did you read the sign about ticks on your way in or out of the trail? *[this question would get at if they have been exposed to the tick drop box more than once]*
2. How old are you?
    a. 18-25
    b. 26-35
    c. 36-45
    d. 46-55
    e. >55
3. Have you ever lived outside of Colorado?
    a. If yes, do you feel comfortable sharing what states/countries you have lived in before?

*Attitudes*

4. How often do you think about ticks or getting tick bites when you are hiking or spending time outdoors? - Rank 1=never, 2= rarely, 3= sometimes, 4=often, 5= always
5. How concerned are you about tick-borne diseases? Rank 1=not concerned, 2= slightly concerned, 3= mildly, 4= moderately, 5= very concerned

*Knowledge*

6. What diseases do ticks in Colorado carry?
7. How do you safely remove a tick that is feeding?
8. *[show a picture] Can you point out the tick or ticks in this picture?"*

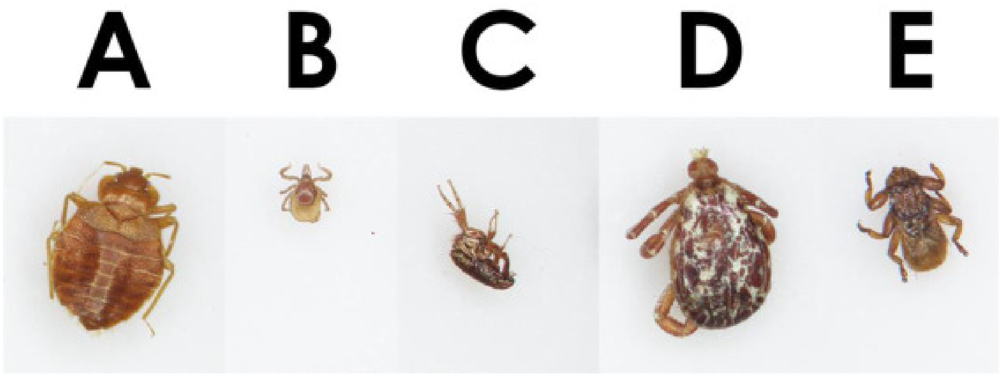


(Cuadera, Mervin Keith Q., et al. 2023)

*Practices*

9. Do you do anything to reduce your chances of getting tick bites? - Open ended questions with tick checks, insect repellents, or avoiding areas with high brush as target outcomes.

### Image Source

Cuadera, Mervin Keith Q., et al. "Knowledge, attitudes, and practices for tick bite prevention and tick control among residents of Long Island, New York, USA." *Ticks and Tick-Borne Diseases* 14.3 (2023): 102124.
