## Supplementary material 3 for "Comparison of tick surveillance approaches: Utilizing trailhead tick-check stations to support tick surveillance and community education"

Table #. Tick knowledge, attitude, and practices (KAP) survey participants' demographics, specifically age group, and current and/or previous residence.

| Survey Topic | Control (N=69) | Intervention (N=30) |
|---|---|---|
| **Age Group** | | |
| **18-25** | 11 (16%) | 1 (3.3%) |
| **26-35** | 19 (28%) | 3 (10%) |
| **36-45** | 7 (10%) | 8 (27%) |
| **46-55** | 13 (19%) | 7 (23%) |
| **>55** | 19 (28%) | 11 (37%) |
| **Current/Previous Residence** | | |
| **Midwest** | 32 (46%) | 8 (27%) |
| **Northeast** | 13 (19%) | 5 (17%) |
| **Pacific** | 27 (39%) | 13 (43%) |
| **South** | 16 (23%) | 10 (33%) |
| **Southeast** | 17 (25%) | 6 (20%) |
| **Mountains** | 4 (5.8%) | 2 (6.7%) |
| **International** | 12 (17%) | 7 (23%) |
| **Outside Colorado** | 62 (90%) | 25 (83%) |
