## Supplementary material 4 for "Comparison of tick surveillance approaches: Utilizing trailhead tick-check stations to support tick surveillance and community education"

Table #. Pearson's correlation values between the responses of different questions from the Knowledge, Attitude, and Practices (KAP) in-person surveys. Each response is characterized under either Variable 1 or Variable 2.

Knowledge, Attitude, and Practices Pearson's Correlation Values

| | Variable 1 | Variable 2 | Pearson's Correlation Value |
|---|---|---|---|
| **Read sign** | Read sign | Read sign | 1 |
| | Correct identification: RMSF | Read sign | 0.269153034140237 |
| | Correct identification: CTF | Read sign | 0.0615429867660051 |
| | Incorrect identification: Lyme disease | Read sign | -0.128944756327002 |
| | Correct practice: tick removal | Read sign | 0.085467146689941 |
| | Correct identification: tick | Read sign | -0.0737209780774486 |
| | Concern: tick bite | Read sign | 0.0809516373350822 |
| | Practices behavior: proactive | Read sign | -0.0219128743686873 |
| | Practices behavior: after | Read sign | 0.0560002159837185 |
| | Practices behavior: proactive and after | Read sign | -0.0428553579981013 |
| | Concern: tick disease | Read sign | -0.00349225795175 |
| **Correct Identification of Rocky Mountain Spotted Fever (RMSF)** | Read sign | Correct identification: RMSF | 0.269153034140237 |
| | Correct identification: RMSF | Correct identification: RMSF | 1 |
| | Correct identification: CTF | Correct identification: RMSF | -0.0901735094004312 |
| | Incorrect identification: Lyme disease | Correct identification: RMSF | -0.305628463384105 |
| | Correct practice: tick removal | Correct identification: RMSF | 0.0454229105915904 |
| | Correct identification: tick | Correct identification: RMSF | 0.110331492464193 |
| | Concern: tick bite | Correct identification: RMSF | 0.0706070030660716 |
| | Practices behavior: proactive | Correct identification: RMSF | 0.0805900077032268 |
| | Practices behavior: after | Correct identification: RMSF | 0.134879204924734 |
| | Practices behavior: proactive and after | Correct identification: RMSF | 0.0416478715212191 |
| | Concern: tick disease | Correct identification: RMSF | 0.0724589235533281 |
| **Correct Identification of Colorado Tick Fever (CTF)** | Read sign | Correct identification: CTF | 0.0615429867660051 |
| | Correct identification: RMSF | Correct identification: CTF | -0.0901735094004312 |
| | Correct identification: CTF | Correct identification: CTF | 1 |
| | Incorrect identification: Lyme disease | Correct identification: CTF | -0.0280799761446314 |
| | Correct practice: tick removal | Correct identification: CTF | -0.0101784189892359 |
| | Correct identification: tick | Correct identification: CTF | -0.0160540324766984 |
| | Concern: tick bite | Correct identification: CTF | 0.0384052405489823 |

| | | | |
|---|---|---|---|
| | Practices behavior: proactive | Correct identification: CTF | -0.0342981152413302 |
| | Practices behavior: after | Correct identification: CTF | 0.0774548984578991 |
| | Practices behavior: proactive and after | Correct identification: CTF | 0.166651853826868 |
| | Concern: tick disease | Correct identification: CTF | -0.0551362684360205 |
| **Incorrect Identification of Lyme Disease** | Read sign | Incorrect identification: Lyme disease | -0.128944756327002 |
| | Correct identification: RMSF | Incorrect identification: Lyme disease | -0.305628463384105 |
| | Correct identification: CTF | Incorrect identification: Lyme disease | -0.0280799761446314 |
| | Incorrect identification: Lyme disease | Incorrect identification: Lyme disease | 1 |
| | Correct practice: tick removal | Incorrect identification: Lyme disease | 0.140910659703795 |
| | Correct identification: tick | Incorrect identification: Lyme disease | 0.00920574617898323 |
| | Concern: tick bite | Incorrect identification: Lyme disease | -0.0951297013477684 |
| | Practices behavior: proactive | Incorrect identification: Lyme disease | 0.0273632234339266 |
| | Practices behavior: after | Incorrect identification: Lyme disease | 0.117178403132897 |
| | Practices behavior: proactive and after | Incorrect identification: Lyme disease | 0.116585580348923 |
| | Concern: tick disease | Incorrect identification: Lyme disease | 0.0103570920642471 |
| **Correct Practice of Proper Tick Removal** | Read sign | Correct practice: tick removal | 0.085467146689941 |
| | Correct identification: RMSF | Correct practice: tick removal | 0.0454229105915904 |
| | Correct identification: CTF | Correct practice: tick removal | -0.0101784189892359 |
| | Incorrect identification: Lyme disease | Correct practice: tick removal | 0.140910659703795 |
| | Correct practice: tick removal | Correct practice: tick removal | 1 |
| | Correct identification: tick | Correct practice: tick removal | 0.104158804734759 |
| | Concern: tick bite | Correct practice: tick removal | 0.137640541156632 |
| | Practices behavior: proactive | Correct practice: tick removal | 0.0559471637132469 |
| | Practices behavior: after | Correct practice: tick removal | 0.0497414937376702 |
| | Practices behavior: proactive and after | Correct practice: tick removal | 0.159835439829598 |
| | Concern: tick disease | Correct practice: tick removal | 0.145342517189329 |
| **Correct Tick Identification** | Read sign | Correct identification: tick | -0.0737209780774486 |
| | Correct identification: RMSF | Correct identification: tick | 0.110331492464193 |
| | Correct identification: CTF | Correct identification: tick | -0.0160540324766984 |
| | Incorrect identification: Lyme disease | Correct identification: tick | 0.00920574617898323 |
| | Correct practice: tick removal | Correct identification: tick | 0.104158804734759 |

| | | | |
|---|---|---|---|
| | Correct identification: tick | Correct identification: tick | 1 |
| | Concern: tick bite | Correct identification: tick | 0.149025337493363 |
| | Practices behavior: proactive | Correct identification: tick | -0.0789545582000777 |
| | Practices behavior: after | Correct identification: tick | 0.0718563137160825 |
| | Practices behavior: proactive and after | Correct identification: tick | 0.020761369963435 |
| | Concern: tick disease | Correct identification: tick | 0.0769783488986253 |
| **Concern for Tick Bites** | Read sign | Concern: tick bite | 0.0809516373350822 |
| | Correct identification: RMSF | Concern: tick bite | 0.0706070030660716 |
| | Correct identification: CTF | Concern: tick bite | 0.0384052405489823 |
| | Incorrect identification: Lyme disease | Concern: tick bite | -0.0951297013477684 |
| | Correct practice: tick removal | Concern: tick bite | 0.137640541156632 |
| | Correct identification: tick | Concern: tick bite | 0.149025337493363 |
| | Concern: tick bite | Concern: tick bite | 1 |
| | Practices behavior: proactive | Concern: tick bite | -0.0127497844855792 |
| | Practices behavior: after | Concern: tick bite | 0.119765127528723 |
| | Practices behavior: proactive and after | Concern: tick bite | 0.0455953498092175 |
| | Concern: tick disease | Concern: tick bite | 0.591235028021589 |
| **Practices Behavior: Proactive** | Read sign | Practices behavior: proactive | -0.0219128743686873 |
| | Correct identification: RMSF | Practices behavior: proactive | 0.0805900077032268 |
| | Correct identification: CTF | Practices behavior: proactive | -0.0342981152413302 |
| | Incorrect identification: Lyme disease | Practices behavior: proactive | 0.0273632234339266 |
| | Correct practice: tick removal | Practices behavior: proactive | 0.0559471637132469 |
| | Correct identification: tick | Practices behavior: proactive | -0.0789545582000777 |
| | Concern: tick bite | Practices behavior: proactive | -0.0127497844855792 |
| | Practices behavior: proactive | Practices behavior: proactive | 1 |
| | Practices behavior: after | Practices behavior: proactive | -0.189748456666312 |
| | Practices behavior: proactive and after | Practices behavior: proactive | 0.293128386529281 |
| | Concern: tick disease | Practices behavior: proactive | 0.139706701192638 |
| **Practices Behavior: After** | Read sign | Practices behavior: after | 0.0560002159837185 |
| | Correct identification: RMSF | Practices behavior: after | 0.134879204924734 |
| | Correct identification: CTF | Practices behavior: after | 0.0774548984578991 |
| | Incorrect identification: Lyme disease | Practices behavior: after | 0.117178403132897 |
| | Correct practice: tick removal | Practices behavior: after | 0.0497414937376702 |
| | Correct identification: tick | Practices behavior: after | 0.0718563137160825 |
| | Concern: tick bite | Practices behavior: after | 0.119765127528723 |

| | | | |
|---|---|---|---|
| | Practices behavior: proactive | Practices behavior: after | -0.189748456666312 |
| | Practices behavior: after | Practices behavior: after | 1 |
| | Practices behavior: proactive and after | Practices behavior: after | 0.622308528391893 |
| | Concern: tick disease | Practices behavior: after | 0.192078703576549 |
| **Practices Behavior: Proactive and After** | Read sign | Practices behavior: proactive and after | -0.0428553579981013 |
| | Correct identification: RMSF | Practices behavior: proactive and after | 0.0416478715212191 |
| | Correct identification: CTF | Practices behavior: proactive and after | 0.166651853826868 |
| | Incorrect identification: Lyme disease | Practices behavior: proactive and after | 0.116585580348923 |
| | Correct practice: tick removal | Practices behavior: proactive and after | 0.159835439829598 |
| | Correct identification: tick | Practices behavior: proactive and after | 0.020761369963435 |
| | Concern: tick bite | Practices behavior: proactive and after | 0.0455953498092175 |
| | Practices behavior: proactive | Practices behavior: proactive and after | 0.293128386529281 |
| | Practices behavior: after | Practices behavior: proactive and after | 0.622308528391893 |
| | Practices behavior: proactive and after | Practices behavior: proactive and after | 1 |
| | Concern: tick disease | Practices behavior: proactive and after | 0.181946188567302 |
| **Concern: Tick Disease** | Read sign | Concern: tick disease | -0.00349225795175 |
| | Correct identification: RMSF | Concern: tick disease | 0.0724589235533281 |
| | Correct identification: CTF | Concern: tick disease | -0.0551362684360205 |
| | Incorrect identification: Lyme disease | Concern: tick disease | 0.0103570920642471 |
| | Correct practice: tick removal | Concern: tick disease | 0.145342517189329 |
| | Correct identification: tick | Concern: tick disease | 0.0769783488986253 |
| | Concern: tick bite | Concern: tick disease | 0.591235028021589 |
| | Practices behavior: proactive | Concern: tick disease | 0.139706701192638 |
| | Practices behavior: after | Concern: tick disease | 0.192078703576549 |
| | Practices behavior: proactive and after | Concern: tick disease | 0.181946188567302 |
| | Concern: tick disease | Concern: tick disease | 1 |
