## Supplementary material 7 for "Comparison of tick surveillance approaches: Utilizing trailhead tick-check stations to support tick surveillance and community education"

Table #. Comparison of tick submissions through CDPHE mail-in system and our (Colorado State University - CSU) tick check stations for Larimer County, CO, during 2024.

| Collection Method | Life Stage* | Species | | Total |
|---|---|---|---|---|
| | | *Dermacentor andersoni* | *Dermacentor variabilis* | |
| By Mail (CDPHE) | | | | |
| | Male | 4 | 1 | 5 |
| | Female | 7 | 14 | 21 |
| | Total | 11 | 15 | 26 |
| By In-person Tick Station (CSU)** | | | | |
| | Male | 29 | 2 | 31 |
| | Female | 43 | 5 | 48 |
| | Total | 72 | 7 | 79 |

*No nymphs or larva were submitted

**Sites included for CSU: CSU Environmental Learning Center (ELC), CSU Mountain Campus, and Soderberg Trail
