## Supplementary material 8 for "Comparison of tick surveillance approaches: Utilizing trailhead tick-check stations to support tick surveillance and community education"

Table #. Analysis from KAP survey results on association, and significance, between participants' previous/current regional residence ("Variable 1") and specific questions asked on survey regarding tick disease identification, tick removal, and tick bite prevention practices ("Variable 2"). Orange highlights what two variable associations had significance ($p \leq 0.05$).

| Variable 1 | Variable 2 | CI 95% | Chi Square | Odds Ratio |
|---|---|---|---|---|
| Read Sign | Correct tick removal & ID | (0.304, 1.74) | 0.464 | 0.728 |
| Read Sign | Correct disease: RMSF | (1.333, 8.756) | 0.008 | 3.381 |
| Read Sign | Correct disease: CTF | (0.059, 92.59) | 0.541 | 2.323 |
| Read Sign | Incorrect disease: Lyme | (0.237, 1.376) | 0.2 | 0.573 |
| Read Sign | Correct tick removal | (0.607, 3.548) | 0.396 | 1.448 |
| Read Sign | Practice behavior: proactive | (0.376, 2.233) | 0.828 | 0.906 |
| Read Sign | Practice behavior: after | (0.486, 3.405) | 0.578 | 1.314 |
| Read Sign | Practice behavior: both | (0.152, 2.866) | 0.67 | 0.766 |
| Midwest | Correct tick removal & ID | (0.427, 2.184) | 0.928 | 0.963 |
| Midwest | Correct disease: RMSF | (0.374, 2.304) | 0.887 | 0.941 |
| Midwest | Correct disease: CTF | (0.038, 58.94) | 0.78 | 1.482 |
| Midwest | Incorrect disease: Lyme | (0.756, 4.118) | 0.187 | 1.737 |
| Midwest | Correct tick removal | (0.579, 2.969) | 0.515 | 1.303 |
| Midwest | Practice behavior: proactive | (0.273, 1.447) | 0.266 | 0.631 |
| Midwest | Practice behavior: after | (1.033, 6.659) | 0.037 | 2.583 |
| Midwest | Practice behavior: both | (0.44, 5.486) | 0.47 | 1.552 |
| Northeast | Correct tick removal & ID | (0.462, 3.948) | 0.6 | 1.31 |
| Northeast | Correct disease: RMSF | (0.81, 7.104) | 0.093 | 2.42 |
| Northeast | Correct disease: CTF | (0.115, 185.126) | 0.239 | 4.602 |
| Northeast | Incorrect disease: Lyme | (0.163, 1.354) | 0.148 | 0.477 |
| Northeast | Correct tick removal | (0.295, 2.406) | 0.74 | 0.843 |
| Northeast | Practice behavior: proactive | (0.787, 9.547) | 0.12 | 2.452 |
| Northeast | Practice behavior: after | (0.471, 4.576) | 0.452 | 1.533 |
| Northeast | Practice behavior: both | (0.604, 9.81) | 0.147 | 2.612 |
| Pacific | Correct tick removal & ID | (0.36, 1.842) | 0.615 | 0.815 |
| Pacific | Correct disease: RMSF | (0.235, 1.537) | 0.293 | 0.619 |
| Pacific | Correct disease: CTF | (0.038, 58.94) | 0.78 | 1.482 |
| Pacific | Incorrect disease: Lyme | (0.38, 1.982) | 0.727 | 0.866 |
| Pacific | Correct tick removal | (0.35, 1.781) | 0.562 | 0.791 |
| Pacific | Practice behavior: proactive | (0.327, 1.726) | 0.489 | 0.75 |
| Pacific | Practice behavior: after | (0.546, 3.435) | 0.487 | 1.375 |

| | | | | |
|---|---|---|---|---|
| Pacific | Practice behavior: both | (0.175, 2.538) | 0.595 | 0.723 |
| South | Correct tick removal & ID | (0.556, 3.581) | 0.475 | 1.385 |
| South | Correct disease: RMSF | (0.909, 6.272) | 0.065 | 2.394 |
| South | Correct disease: CTF | | | |
| South | Incorrect disease: Lyme | (0.235, 1.478) | 0.246 | 0.591 |
| South | Correct tick removal | (0.505, 3.173) | 0.621 | 1.25 |
| South | Practice behavior: proactive | (0.259, 1.637) | 0.343 | 0.648 |
| South | Practice behavior: after | (0.482, 3.642) | 0.544 | 1.362 |
| South | Practice behavior: both | (0.075, 2.366) | 0.421 | 0.557 |
| Southwest | Correct tick removal & ID | (0.157, 1.103) | 0.071 | 0.426 |
| Southwest | Correct disease: RMSF | (0.281, 2.46) | 0.79 | 0.879 |
| Southwest | Correct disease: CTF | | | |
| Southwest | Incorrect disease: Lyme | (0.411, 2.883) | 0.888 | 1.067 |
| Southwest | Correct tick removal | (0.225, 1.535) | 0.27 | 0.596 |
| Southwest | Practice behavior: proactive | (0.582, 4.518) | 0.371 | 1.553 |
| Southwest | Practice behavior: after | (0.218, 2.167) | 0.574 | 0.743 |
| Southwest | Practice behavior: both | (0.225, 4.385) | 0.878 | 1.148 |
| Mountains | Correct tick removal & ID | (0.562, 104.504) | 0.158 | 3.842 |
| Mountains | Correct disease: RMSF | (0.437, 16.575) | 0.223 | 2.688 |
| Mountains | Correct disease: CTF | | | |
| Mountains | Incorrect disease: Lyme | (0.233, 11.215) | 0.716 | 1.333 |
| Mountains | Correct tick removal | (0.052, 2.456) | 0.306 | 0.43 |
| Mountains | Practice behavior: proactive | (0.213, 10.269) | 0.793 | 1.22 |
| Mountains | Practice behavior: after | (0.175, 8.59) | 0.685 | 1.48 |
| Mountains | Practice behavior: both | (0.058, 12.04) | 0.725 | 1.63 |
| International | Correct tick removal & ID | (0.311, 2.431) | 0.776 | 0.865 |
| International | Correct disease: RMSF | (0.383, 3.58) | 0.723 | 1.227 |
| International | Correct disease: CTF | | | |
| International | Incorrect disease: Lyme | (0.253, 1.988) | 0.492 | 0.705 |
| International | Correct tick removal | (0.579, 4.806) | 0.35 | 1.611 |
| International | Practice behavior: proactive | (0.384, 3.222) | 0.878 | 1.076 |
| International | Practice behavior: after | (0.041, 1.154) | 0.083 | 0.294 |
| International | Practice behavior: both | (0.114, 3.791) | 0.813 | 0.869 |
