## Supplementary material 6 for "Comparison of tick surveillance approaches: Utilizing trailhead tick-check stations to support tick surveillance and community education"

### Passive Tick Surveillence Comparison 2024: Mail-in System v. Tick Check Stations

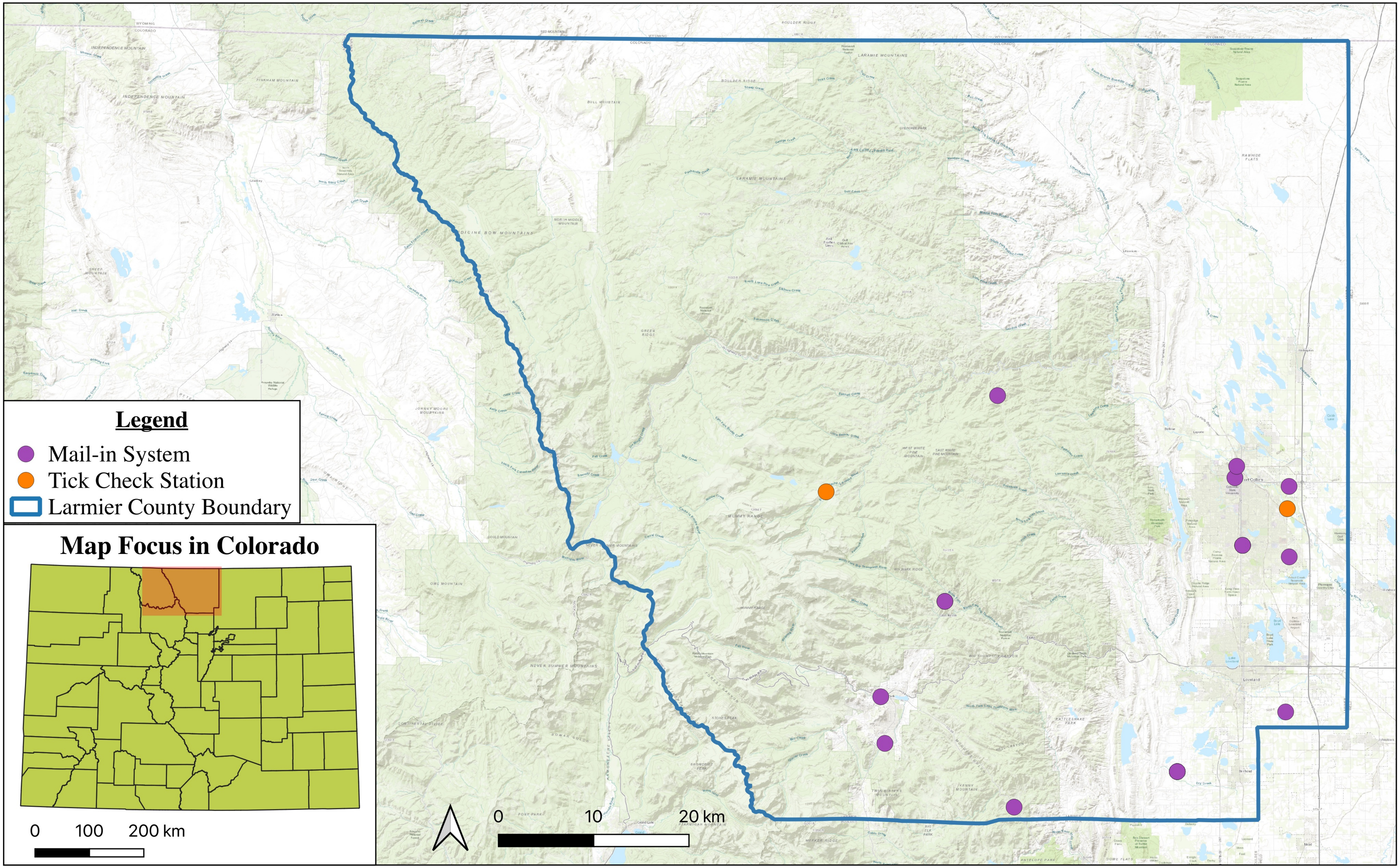


Supplement Figure #. Comparision of location of ticks collected from passive surveillence either via mail-in system (CDPHE) or tick check station (CSU) in Larmier County, CO, during 2024.
